# Dynamic posttranslational modifications of cytoskeletal proteins unveil hot spots under nitroxidative stress

**DOI:** 10.1101/2021.01.24.427971

**Authors:** Eva Griesser, Venukumar Vemula, Andreia Mónico, Dolores Pérez-Sala, Maria Fedorova

**Author notes:** Corresponding authors: Dr. Maria Fedorova, Institut für Bioanalytische Chemie, Biotechnologisch-Biomedizinisches Zentrum, Deutscher Platz 5, 04103 Leipzig, Germany.; Dr. Dolores Pérez-Sala, Centro de Investigaciones Biológicas Margarita Salas, C.S.I.C., Ramiro de Maeztu, 9, 28040 Madrid, Spain. Present address: Drug Discovery Sciences, Boehringer Ingelheim Pharma GmbH & Co. KG, Biberach an der Riss, Germany. Present address: Linnaeus University, Dept. of Chemistry and Biomedical Sciences, Universitetskajen, 391 82 Kalmar, Sweden (Venukumar Vemula).

## Abstract

The cytoskeleton is a supramolecular structure consisting of interacting protein networks that support cell dynamics in essential processes such as migration and division, as well as in responses to stress. Fast cytoskeletal remodeling is achieved with the participation of regulatory proteins and posttranslational modifications (PTMs). Redox-related PTMs are emerging as critical players in cytoskeletal regulation. Here we used a cellular model of mild nitroxidative stress in which a peroxynitrite donor induced transient changes in the organization of three key cytoskeletal proteins, i.e., vimentin, actin and tubulin. Nitroxidative stress-induced reconfiguration of intermediate filaments, microtubules and actin structures were further correlated with their PTM profiles and dynamics of the PTM landscape. Using high-resolution mass spectrometry, 62 different PTMs were identified and relatively quantified in vimentin, actin and tubulin proteins, including 12 enzymatic, 13 oxidative and 2 nitric oxide-derived modifications as well as 35 modifications by carbonylated lipid peroxidation products, thus evidencing the occurrence of a chain reaction with formation of reactive species and the activation of multiple signaling pathways. Our results unveil the presence of certain modifications under basal conditions and their modulation in response to stress in a target-, residue- and reactive species-dependent manner. Moreover, we identified protein PTM “hot spots”, such as the single cysteine residue of vimentin, supporting its role in PTM cross-talk and redox sensing. Finally, identification of novel PTMs in these proteins may pave the way for unveiling novel cytoskeleton regulatory mechanisms.

## Introduction

The cytoskeleton is a self-organizing supramolecular structure constituted by a hierarchically ordered system of a large number of mutually interdependent molecules. It provides a physical and biochemical connection between the cell and its extracellular environment by receiving, integrating, and transducing extra- and intracellular stimuli, and generating coordinated forces that allow cells to move, change shape and divide (1–3). Microtubules, intermediate filaments and actin are the three main components of the cytoskeleton (1,4). Their constituent proteins are capable of self-assembly into high dynamic polymers, co-existing in equilibrium with oligomeric pools, and undergoing constant exchange in the cell in response to external or internal cues. Regulation of cytoskeletal dynamics is crucial since cellular responses need to be fast and versatile. Cytoskeletal remodeling relies mainly on the participation of regulatory and crosslinker proteins and molecular motors (5,6). For instance, over 150 cellular proteins containing actin-binding domains, and thus, capable to interact with and/or regulate actin dynamics, have been reported by now (7). Posttranslational modifications (PTMs) are crucial for the rapid modulation of protein function. Phosphorylation/dephosphorylation events constitute a central signaling mechanism and are key for the spatio-temporal regulation of the assembly or interactions of cytoskeletal structures (8,9). Other enzymatic PTMs important for cytoskeletal modulation include acetylation, polyubiquitination and SUMOylation, glycosylation or proteolytic cleavage (9–11). In addition, non-enzymatic PTMs, in particular those induced by oxidants and electrophiles, are emerging as fast and versatile mechanisms for cytoskeletal regulation (12,13). Redox balance is a basic mechanism of cellular homeostasis, achieved by the equilibrium between a variety of oxidants, often reactive oxygen and nitrogen species (ROS/RNS), and the cellular antioxidant defenses. ROS/RNS transduce the information via oxidative modifications of biomolecules including nucleic acids, proteins, lipids and metabolites (14,15). Oxidative protein PTMs are one of the main mechanisms of redox regulation among which thiol-based redox reaction probably is the most studied one so far (16). Moreover, in many cases, reversible oxidative modifications can act as “redox switches”, much like phosphorylation or other enzymatic modifications. In addition, through their actions on lipids, ROS/RNS can give rise to a plethora of structurally diverse reactive lipid peroxidation products (LPPs) that can form adducts with proteins in a process called lipoxidation (17,18). These modifications can contribute to signaling and/or oxidative protein damage under oxidative stress conditions, and affect residues involved in redox signaling, for which we will use the term (lip)oxidative modifications. Cys-modifications are often involved in redox sensing via sulfenation, S-glutathionylation, or S-nitrosation (frequently termed nitrosylation), among others (16). Furthermore, Cys residues are frequently modified by reactive LPPs (19). Lys residues are susceptible to oxidation (protein carbonyl formation) and modifications by LPPs (19). Interestingly, Cys- and Lys-residues are often involved in protein-protein or protein-DNA interactions, and in regulating a large number of signaling pathways via several other PTMs (20,21). In turn, Tyr residues can be involved in redox switches through oxidation and nitration and play an important role in cross-talk with the well-studied Tyr phosphorylation (22,23).

Certain amino acid residues in given proteins represent “hot spots” for PTMs because they are susceptible to many types of PTMs, including redox regulated ones, and are frequently modified to a larger extent than other residues (24,25). Hot spot residues are often conserved in proteins and play regulatory roles in response to various stimuli, thus contributing to PTM cross-talk. Indeed, the cross-talk between PTMs allows to formulate the hypothesis of "PTM codes", which represents an additional level of cell regulation and signaling complexity (26), with the histone code phenomena being a well-known example (27). However, the role of (lip)oxidative modifications in PTM cross-talks is still not completely understood, mostly due to the analytical challenges in their high-throughput detection and quantification.

Cytoskeletal remodeling in response to redox changes, as well as the importance of redox-related modifications of cytoskeletal proteins in essential processes such as migration, division and response to stress, and its interplay with other PTMs require in-depth profiling of cytoskeletal PTM codes. In a previous study we established a dynamic cellular model based on rat primary cardiac cells treated with a peroxynitrite donor (3-morpholinosydnonimine, SIN-1) for various time periods. Initial proteomic analysis showed the importance of cytoskeletal proteins as targets for modification by lipid peroxidation products (28). Here we have used this model to track the dynamics of cytoskeletal rearrangements and relatively quantify changes of PTM profiles for selected cytoskeletal proteins. Thus, we have combined confocal microscopy evaluation of morphological changes in the three main cytoskeletal systems: microtubules, intermediate filaments and microfilaments, upon mild nitroxidative stress with a detailed mapping of over 60 different PTMs in protein building blocks of these systems, namely, α and β tubulin, vimentin and actin, using high resolution tandem mass spectrometry. Using hierarchical clustering and principal component analysis (PCA), we unveil a cascade of enzymatic and oxidative PTMs constituting different dynamic PTM patterns and identify several amino acid residues representing “hot spots” for PTM occurrence and crosstalk.

## Material and methods

### Materials

Dulbecco’s Modified Eagle Medium/Ham’s F-12 (DMEM/F12), phosphate buffered saline (PBS), fetal bovine serum (FBS), penicillin-streptomycin, L-glutamine, non-essential amino acids, sodium pyruvate, and gelatin were obtained from Life Technologies GmbH (Darmstadt, Germany). Horse serum, trypsin-EDTA solution, paraformaldehyde (PFA), triton-X-100, bovine serum albumin (BSA), 4’,6-diamidino-2-phenylindole (DAPI), thiourea, ß-mercaptoethanol, Na-deoxycholate, Nonidet P-40, formic acid, primary mouse monoclonal anti-β-tubulin antibody (T8328) and all ammonium and sodium salts were purchased from Sigma-Aldrich GmbH (Taufkirchen, Germany). 3-morpholinosydnonimine (SIN-1) was purchased from Enzo Life Sciences GmbH (Lörrach, Germany). Urea, SDS, glycerol and Tris(2-carboxyethyl)phosphine hydrochloride (TCEP) were obtained from Carl Roth GmbH + Co. KG (Karlsruhe, Germany). CHAPS, bromophenol blue, glycine, Coomassie Brilliant Blue G-250 and trypsin were purchased from Serva Electrophoresis GmbH (Heidelberg, Germany). Tris, iodoacetamide (IAA) and EDTA were from Applichem GmbH (Darmstadt, Germany). Acetonitrile (ULC-MS grade) was from Biosolve (Valkenswaard, Netherlands) and acetone (LiChroSolv) from Merck Chemicals GmbH (Darmstadt, Germany). Agarose and Alexa488-conjugated anti-vimentin V9 antibodies (sc-6260) were obtained from Santa Cruz Biotechnology, Inc. (Heidelberg, Germany), phalloidin-Alexa568 and Alexa488-conjugated anti-mouse antibody were from Molecular Probes (Life Technologies GmbH).

### Cell culture

Primary rat cardiac cells (Innoprot, Elexalde Derio, Spain) were cultured on gelatin-coated 75 cm^2^ flasks (CELLSTAR, Greiner Bio-One GmbH, Frickenhausen, Germany) in DMEM/F12 medium supplemented with 20% FBS, 5% horse serum, 2 mmol/L L-glutamine, 3 mmol/L sodium pyruvate, 0.1 mmol/L non-essential amino acids, 100 U/mL penicillin and 100 μg/mL streptomycin at 37 °C (humidified atmosphere of 5% CO_2_ and 95% air). When cells reached 80% confluence and 24 h before treatment medium was replaced by serum-free medium (DMEM/F12 supplemented with 100 U/mL penicillin and 100 μg/mL streptomycin). Cells (passages 3-5) were treated with 10 μmol/L SIN-1 for 15 min, 30 min, 70 min and 16 h.

### Immunofluorescence of cytoskeletal proteins actin, vimentin and tubulin

Cells were grown on gelatin-coated cover slips in 24-well-plates, treated with SIN-1, washed twice with PBS, fixed with PFA (4% w/v in PBS, 30 min, RT) and washed with PBS three more times. Cells were permeabilized with 0.1% (v/v) Triton-X-100 in PBS (20 min, RT), washed twice with PBS, blocked with 1% BSA (w/v) in PBS (blocking solution, 1 h) and incubated with Alexa488-conjugated anti-vimentin V9 antibody, primary mouse monoclonal anti-β-tubulin antibody (both 1:200 in blocking solution, 1 h), or phalloidin-Alexa568 (1:40 in blocking solution, 30 min). For tubulin visualisation, Alexa488-conjugated secondary anti-mouse antibody (1:200 in blocking solution, 1 h) was used. After incubations cells were washed with PBS (three times). For co-visualization of actin and vimentin cells were first incubated with anti-vimentin-Alexa488 followed by phalloidin-Alexa568. Nuclei were counterstained with DAPI (3.3 μg/mL in PBS, 15 min) followed by washes (two times PBS, one time water). Cover slips were mounted on glass slides using FluorSave (Calbiochem, Merck Millipore). Cells were visualized with a confocal laser scanning microscope (Leica TCS SP5, Leica Microsistemas S.L.U., Barcelona, Spain. Individual sections were taken every 0.5 μm with a 63× objective (NA 1.4, oil) and single sections or overall projections are shown, as indicated. Images were acquired using Leica Application Suite (LAS AF) software. Image analysis was performed with FIJI ImageJ software (29).

### Protein extraction

After SIN-1 treatment cells were scraped in ice-cold PBS and collected by centrifugation (10 min, 1000 × *g*, 4 °C). Cell pellets were washed with ice-cold PBS (two times) and resuspended in urea lysis buffer (7 mol/L urea, 2 mol/L thiourea, 2% CHAPS in 50 mmol/L Tris-HCl, pH 7.5). Samples were sonicated on ice using a Vibra-Cell tip sonicator (20 kHz, 1 min with on/off pulses of 5 s each, 40% amplitude; Sonics & Materials, Inc., Newtown, CT, USA) and centrifuged (20 min, 10,000 × *g*, 4 °C). Supernatants were collected and reversibly modified cysteines were reduced with 5 mmol/L TCEP (60 min, 37 °C, 350 rpm) followed by their alkylation with 20 mmol/L iodoacetamide (1 h, 27 °C, 350 rpm, in the dark). Proteins were precipitated using acetone (five volumes, −20 °C, overnight). Samples were centrifuged (30 min, 10,000 × *g*, RT), pellets were washed with 1 mL of ice-cold acetone, dried by vacuum concentration and redissolved in urea lysis buffer. Protein concentrations were determined by Bradford assay.

### Immunoprecipitation of vimentin

Proteins (200 μg in urea lysis buffer) were diluted in RIPA buffer (total 960 μL; 150 mmol/L NaCl, 1 mmol/L EDTA, 0.1% SDS, 1% Na-deoxycholate, 1% Nonidet P-40, 50 mmol/L Tris-HCl pH 7.3) and incubated with 40 μL of agarose-conjugated anti-vimentin V9 antibody on a rotary shaker overnight at 4 °C. Samples were centrifuged (1 min, 14,000 × *g*, 4 °C) and washed with ice-cold PBS (four times). Proteins were eluted with 25 μL Laemmli sample buffer (62.5 mmol/L Tris-HCl pH 6.8, 20% v/v glycerol, 2% w/v SDS, 5% v/v ß-mercaptoethanol, 0.01% w/v bromophenol blue; 95 °C, 5 min) and separated by SDS-PAGE.

### Gel electrophoresis

Proteins (25 μg) were mixed with Laemmli sample buffer, separated by SDS-PAGE (12% T, 1 mm, 200 V; BioRad mini protean III cell; BioRad Laboratories GmbH, München, Germany) and stained with Coomassie Brilliant Blue G-250.

### In-gel tryptic digestion

For analysis of the three cytoskeletal structures studied, different enrichment strategies were utilized. Vimentin was enriched by immunoprecipitation, separated by SDS-PAGE, and digested with trypsin. For the analysis of actin and tubulin proteins the whole cell extract was separated by SDS-PAGE. Protein bands corresponding to actin, tubulin and vimentin were excised (as indicated in Figure S1), destained with 50% (v/v) acetonitrile in 50 mmol/L NH_4_HCO_3_ (1 h, 37 °C, 750 rpm), dehydrated with 100% acetonitrile and dried under vacuum. Proteins were digested with trypsin in 3 mmol/L NH_4_HCO_3_ (250 ng, 4 h, 37 °C, 550 rpm) and peptides were extracted by consecutive incubations with 100%, 50% (v/v; aqueous solution) and 100% acetonitrile (15 min sonication for each step). Combined extracts were vacuum concentrated. Peptides were dissolved in 10 μL (tubulin, vimentin) or 20 μL (actin) of 0.5% formic acid in 60% aqueous acetonitrile and further diluted 1:10 with 0.1% formic acid in 3% aqueous acetonitrile.

### LC-MS/MS

A nano-Acquity UPLC (Waters GmbH, Eschborn, Germany) was coupled online to an LTQ Orbitrap XL ETD mass spectrometer equipped with a nano-ESI source (Thermo Fischer Scientific, Bremen, Germany). Eluent A was aqueous formic acid (0.1% v/v), and eluent B was formic acid (0.1% v/v) in acetonitrile. Samples (10 μL) were loaded onto the trap column (nanoAcquity symmetry C18, internal diameter 180 μm, length 20 mm, particle diameter 5 μm) at a flow rate of 10 μL/min. Peptides were separated on a BEH 130 column (C18-phase, internal diameter 75 μm, length 100 mm, particle diameter 1.7 μm) with a flow rate of 0.4 μL/min. Peptides from vimentin and tubulin samples were separated using several linear gradients from 3% to 9% (2.1 min), 9.9% (1.9 min), 17.1% (10 min), 18% (0.5 min); 20.7% (0.2 min), 22.5% (3.1 min), 25.6% (3 min), 30.6% (5 min), 37.8% (2.8 min) and finally to 81% eluent B (2 min). Together with an equilibration time of 12 min the samples were injected every 46 min. Peptides from actin samples were separated using two step gradients from 3 to 35% eluent B over 90 min and then to 85% eluent B over 4 min. After 5 min at 85% eluent B the column was equilibrated for 15 min and samples were injected every 120 min. The transfer capillary temperature was set to 200 °C and the tube lens voltage to 120 V. An ion spray voltage of 1.5 kV was applied to a PicoTip online nano-ESI emitter (New Objective, Berlin, Germany). The precursor ion survey scans were acquired at an orbitrap (resolution of 60,000 at *m/z* 400) for a *m/z* range from 400 to 2000. CID-tandem mass spectra (isolation width 2, activation Q 0.25, normalized collision energy 35%, activation time 30 ms) were recorded in the linear ion trap by data-dependent acquisition (DDA) for the top six most abundant ions in each survey scan with dynamic exclusion for 60 s using Xcalibur software (version 2.0.7). The first set of LC-MS/MS was used to identify highly abundant unoxidized peptides using the Sequest search engine (Proteome Discoverer 1.4, Thermo Scientific) against the Uniprot *Rattus Norvegicus* database (downloaded on March, 11 2014), allowing up to two missed cleavages and a mass tolerance of 10 ppm for precursor ions and 0.8 Da for product ions. Oxidation of methionine and carbamidomethylation of cysteine were used as variable modifications and results were filtered for rank 1 peptides and charge-dependent scores (Xcorr ≥ 2.0, 2.25, 2.5, and 2.75 for charge states 2, 3, 4, and 5). The list of *m/z* values and retention times (±1 min) corresponding to identified peptides was exported and used as exclusion list of unmodified peptides for the second LC-MS/MS analysis, which was performed by DDA using CID and ETD (activation time 100 ms, isolation width 2 u) fragmentation techniques (Figure S2).

### Database search

The acquired tandem mass spectra were searched against the Uniprot *Rattus Norvegicus* database using Sequest search engine (Proteome Discoverer 1.4, Thermo Scientific), allowing up to two missed cleavages and a mass tolerance of 10 ppm for precursor ions and 0.8 Da for product ions. Oxidation of methionine and carbamidomethylation of cysteine were used as variable modifications. Only protein identifications with a protein score ≥ 10 and minimum 2 peptides per protein with peptides ranked on position 1 and charge-dependent scores (Xcorr ≥ 2.0, 2.25, 2.5, and 2.75 for charge states 2, 3, 4, and 5) were considered. Accession numbers of the identified proteins from all conditions were used to create limited FASTA files for each actin, tubulin and vimentin, which were used for the identification of their modifications. Additionally, not identified actin and tubulin isoforms were added. Database search was performed using 62 different PTMs as variable modifications (Table S1), including 12 enzymatic, 13 oxidative and 2 NO-based modifications as well as 35 modifications by carbonylated lipid peroxidation products (oxoLPPs), which we recently identified in SIN-1 treated cardiac cells (28). Analyzed PTMs covered eleven amino acid residues including Lys (48 modifications), Arg (thirty), Cys (seventeen), His (thirteen), Trp (eight), Tyr (five), Phe (four), Pro (three), Thr (two), Met (two), and Ser (one). For each search, oxidation of Cys, Met and Trp, and carbamidomethylation were always used as variable modifications in addition to a set of three other modifications. Only peptides with medium and high confidence, charge-dependent scores as indicated above and ranked on position 1 were considered. All modified peptides and their modification sites were validated by manual inspection of their MS/MS spectra.

Additionally, modified peptides from two independent experiments (phosphopeptide enrichment using TiO_2_ and selective reduction of nitrosation and glutathionylation followed by iodoTMTsixplex labeling) were included into the final results (see Supplementary Information).

### Label-free relative quantification

Label-free relative quantification was performed using Progenesis QI for proteomics (Nonlinear Dynamics, Newcastle, UK) using LC-MS data from three biological replicates analysed in three technical replicates. Only peptides showing regulation with ANOVA p-value <0.05 were considered for further analysis. The resulting peptide list from Progenesis QI analysis was compared with the list of identified modified peptides. Results were evaluated by principal component analysis (PCA) using EZinfo 2.0 within the MassLynx software (version 4.1; Waters GmbH, Eschborn, Germany). Hierarchical clustering of modification sites was performed using GENESIS software (version 1.8.1) (30).

## Results

### Mild nitroxidative stress induces transient remodeling of cytoskeletal structures

Induction of nitroxidative stress by treatment of a primary cardiac cell culture with 10 μmol/L SIN-1 followed by immunocytochemistry and confocal microscopy analysis, revealed marked rearrangements of intermediate filaments (vimentin), microfilaments (actin) and microtubules (tubulin) (Figure 1). In non-treated (control) cells, vimentin appeared as a homogeneously extended filament network, with occasional perinuclear bundles (17% of the cells) (Figure 1A, left panels). Treatment with SIN-1 elicited a marked retraction of vimentin filaments from the cell periphery and increased condensation in the perinuclear area. This effect was quantitated in single cells by measuring the area of the cell occupied by vimentin filaments with respect to that of actin staining, as an index of total cell area (Figure 1B). This revealed an approximately 20% vimentin network retraction after 15 min treatment. This effect persisted at least during 70 min, but 16 h after addition of SIN-1, the normal morphology was eventually recovered.

**Figure 1:**
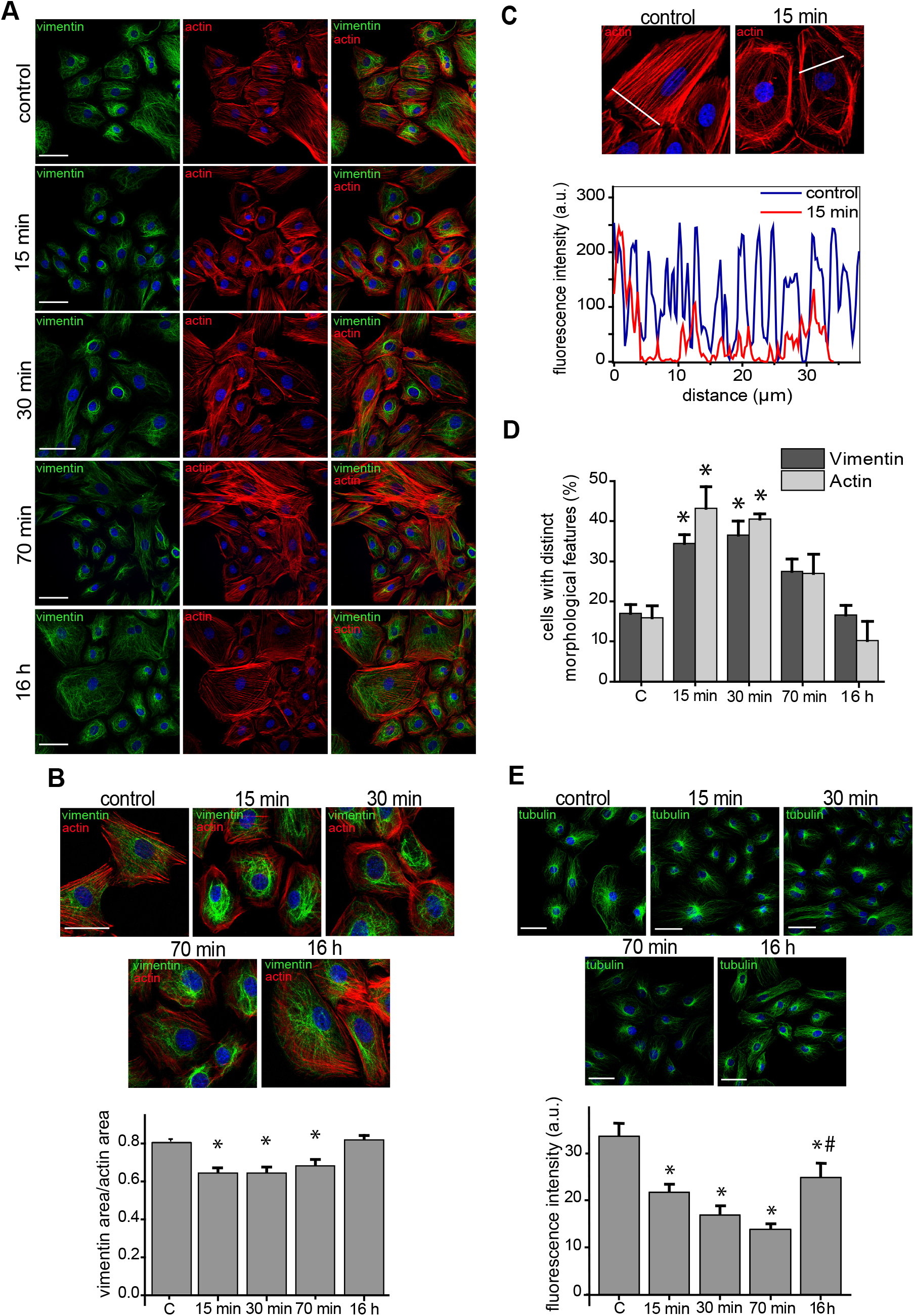
Morphological changes of actin, vimentin and tubulin induced by nitroxidative stress. (A) Rat primary cardiac cells were treated with SIN-1 (10 μmol/L) for the indicated times and the distribution of f-actin (red) and vimentin (green), was assessed by phalloidin staining and immunofluorescence, respectively. Nuclei were counter-stained with DAPI (blue) in all panels. (B) The distribution of f-actin and vimentin are shown in more detail. Images are representative of three independent experiments. Scale bars, 50 μm. The cellular areas covered by the vimentin and the actin cytoskeletons were measured by FIJI and their ratios calculated for at least 20 cells per condition. Data are shown in the lower histogram as mean ± SEM (* p<0.005 vs. control, depicted as “C” in the graph). (C) The distribution of f-actin was assessed as in (A) and the fluorescence intensity profiles along the white lines were obtained with FIJI and are depicted in the lower panel as a function of the distance. (D) Proportion of cells showing distinct morphological features, characteristic of nitroxidative stress, as detailed in the text, namely, perinuclear condensation for vimentin and loss of stress fibers for actin. Data are shown as mean ± SEM (*p<0.01 vs. control) of 40-50 cells per condition, obtained from three independent experiments. (E) Cells were treated with SIN-1, as indicated, and the distribution of β-tubulin was monitored by immunofluorescence. The lower panel represents the density of the microtubule network at the cell periphery estimated from the fluorescent intensity of an area adjacent to cell membrane. Graphed values are mean ± SEM of at least 20 cells per time point (*p<0.05 with respect to control, depicted as “C” in the graph, #p<0.05 with respect to the 70 min time point); a.u., arbitrary units.

Staining of actin showed that most untreated cells (>80%) displayed robust stress fibers that frequently spanned the whole cell (Figure 1A, middle panels). Treatment with SIN-1 resulted in a marked decrease in the abundance of stress fibers and an increase in the proportion of dot-like structures. Additionally, SIN-1-treated cells showed increased staining of actin along the plasma membrane (cortical actin) and reduced or more diffuse cytoplasmic staining (Figure 1A and C). This morphological shift was clearly evidenced in single cells by comparing the actin fluorescence intensity profiles across the cell body (Figure 1C). The proportions of cells showing predominant perinuclear vimentin distribution and predominant cortical actin vs stress fibers for the different time points are shown in Figure 1D. Intense perinuclear vimentin condensation was observed in over 35% of the cells at 15 and 30 min after SIN-1 treatment, whereas over 40% of the cells displayed decreased stress fibers and increased cortical actin at these time points. However, these effects were attenuated after 70 min, and 16 h after SIN-1 addition cells had recovered nearly normal appearance according to these parameters (Figure 1D).

The organization of microtubules in control cells showed an extended distribution from the juxtanuclear area to the cell periphery. At 15, 30 and 70 min after SIN-1 addition, microtubules showed an increased juxtanuclear condensation, with a concomitant loss of peripheral density, quantitated as the intensity of fluorescence at the cell periphery (Figure 1E). As observed for vimentin and actin, 16 h after the beginning of the treatment the morphology of microtubules was restored.

### Highly conserved vimentin residues constitute PTM hot pots

A bottom-up proteomic approach was used to identify possible PTMs accompanying morphological changes induced by nitroxidative stress, focusing on three of the main cytoskeletal proteins, namely, vimentin, actin and tubulin. In addition, availability of this dynamic cellular model of nitroxidative stress allowed performing relative label-free or TMT-based (in the case of Cys modifications) quantification of the modified peptides. A detailed overview of all modifications and their quantitative analysis is presented as supplementary information, and the most relevant findings will be described in this section. A plethora of PTMs was detected on vimentin during the treatment with SIN-1. Overall, 225 modified peptides were detected, highlighting 61 modification sites (Table 1 and S2, Figure S5). Interestingly, most modifications were dynamic, and their nature and/or extent varied with treatment time. The 61 vimentin modification sites showed different accessibility to PTMs. Most of them underwent one or two modification types; however several residues were identified as PTM “hot spots” (Figure 2A). The single Cys residue of vimentin, Cys328, which is conserved in all type III intermediate filament proteins, has been reported to be the target of diverse PTMs under various pathophysiological conditions and experimental models (see (13) for review). Here, Cys328 was observed to accommodate at least seven different modifications, including dioxidation and trioxidation (formation of sulfinic and sulfonic acids, respectively), modification by the reactive aldehydes glyoxal, hexenal and pentenal, and the well-known modifications S-glutathionylation and S-nitrosation. Thus, from our results, this Cys residue emerges as a hot spot. Interestingly, the proportion of non-modified Cys328 transiently increased shortly after addition of SIN-1, suggesting the existence of some basal modifications that could be reversed upon stress. Indeed, Cys328 glutathionylation was detected under basal conditions and transiently decreased upon SIN-1 addition, although it increased at later time-points (i.e. 30 and 70 min after SIN-1 addition), and returned to basal levels after 16 h (Figure 2C, Table S3). In contrast, Cys328 nitrosation and sulfonic acid formation suffered a moderate and sustained increase. Vimentin also possesses a single Trp residue (Trp290) that is conserved between species as well as in other members of the type III intermediate filament family. Trp290 underwent at least six oxidative modifications, including oxidation, dioxidation and trioxidation, formation of kynurenine, hydroxkynurenine and oxolactone. These modifications reached a maximum level 30 min after SIN-1 addition and decayed afterwards (Figure 2C). Importantly, Lys residues were the most frequently modified with 20 out of 22 available Lys residues modified by 12 different PTM types, including formylation (18 positions), oxoLPP adducts (hexenal - sixteen; hydroxy-nonenal (HNE)-three, methylglyoxal/malondialdehyde (MDA) – four, glyoxal, hydroxy-hexenal (HHE), oxo-pentanal, and pentenal – each at one position), dimethylation (three), acetylation (two), methylation (one), and succinylation (one). Of Lys residues, Lys129 emerged as a hot spot with seven PTMs including enzymatic acetylation, succinylation, four oxoLPP adducts, and formylation. For other Lys residues, namely Lys223, Lys235, and Lys236, four PTMs were detected. Remarkably, the corresponding unmodified peptides covering those residues showed a transient increase during the treatment, suggesting the reversal of some basal modification(s). Modification by methylglyoxal/MDA appeared as transient events, whereas formylation and HNE modification (in the case of Lys235) accumulated during treatment. Tyr residues were oxidized to dihydroxyphenylalanine (DOPA, Tyr38, 53, 61, 150, 291 and 400), dopaquinone (DQ, Tyr38, 53, 61, 291), and trihydroxyphenylalanine (TOPA, Tyr53, 61 and 291). Among them Tyr61 emerged as a hot spot showing four PTMs, including three oxidations and nitration. Of other residues, His was modified by hexenal (His238, 253, 437, 461, and 462), HNE (His437), and oxidation (His461 and 462). Modifications of Met to sulfoxide (ten out of eleven available positions) and sulfone (six residues) were the most abundant oxidative PTMs. Formation of protein carbonyls on vimentin via direct oxidation of amino acid residues was shown for Thr327, 426, 458 (2-amino-3-ketobutyric acid) and Arg158 (glutamic semialdehyde). Pro 57 was transformed into pyroglutamic acid and mono- and dihydroxylated. Phosphorylation is a key modification in vimentin, since it is involved in the disassembly of filaments. Multiple phosphorylation sites were detected during treatment with SIN-1, including Thr37 and Thr458, Tyr38, and ten Ser residues (positions 25, 39, 49, 56, 66, 83, 144, 226, 430, and 459). Thus, most phosphorylation events affected vimentin head domain. Detailed quantitative data on the modification of vimentin residues, analyzed by hierarchical clustering are presented in Figure S7A, and principal component analysis is presented in Figure S8. In summary, almost all quantifiable Lys formylation sites, as well as oxidative modifications, represented mainly by Met, Pro, and Tyr mono- and dioxidation were significantly upregulated with a maximum at 16 h after SIN-1 treatment. Another set of peptides containing oxidized Trp or Met residues showed their maximum 30 min or 70 min after SIN-1 treatment. However, other modifications, and in particular those related to lipoxidation, showed a transient pattern, with the exception of HNE-modified Lys235. PCA analysis (Figure S8) allowed specifying several clusters of closely projected PTMs. Thus, most of the oxidative modifications as well as Lys formylation sites showed similar direction and length of the PTM vectors. Interestingly, several residues identified as modification “hot spots” (Figure 2A) showed a different behavior than most of the oxidation and formylation sites – thus Cys328 and Trp290 had close projections, whereas oxoLPP modifications of Lys129 and 223/235/236 formed a different cluster together with oxidized His461/462.

**Table 1:**
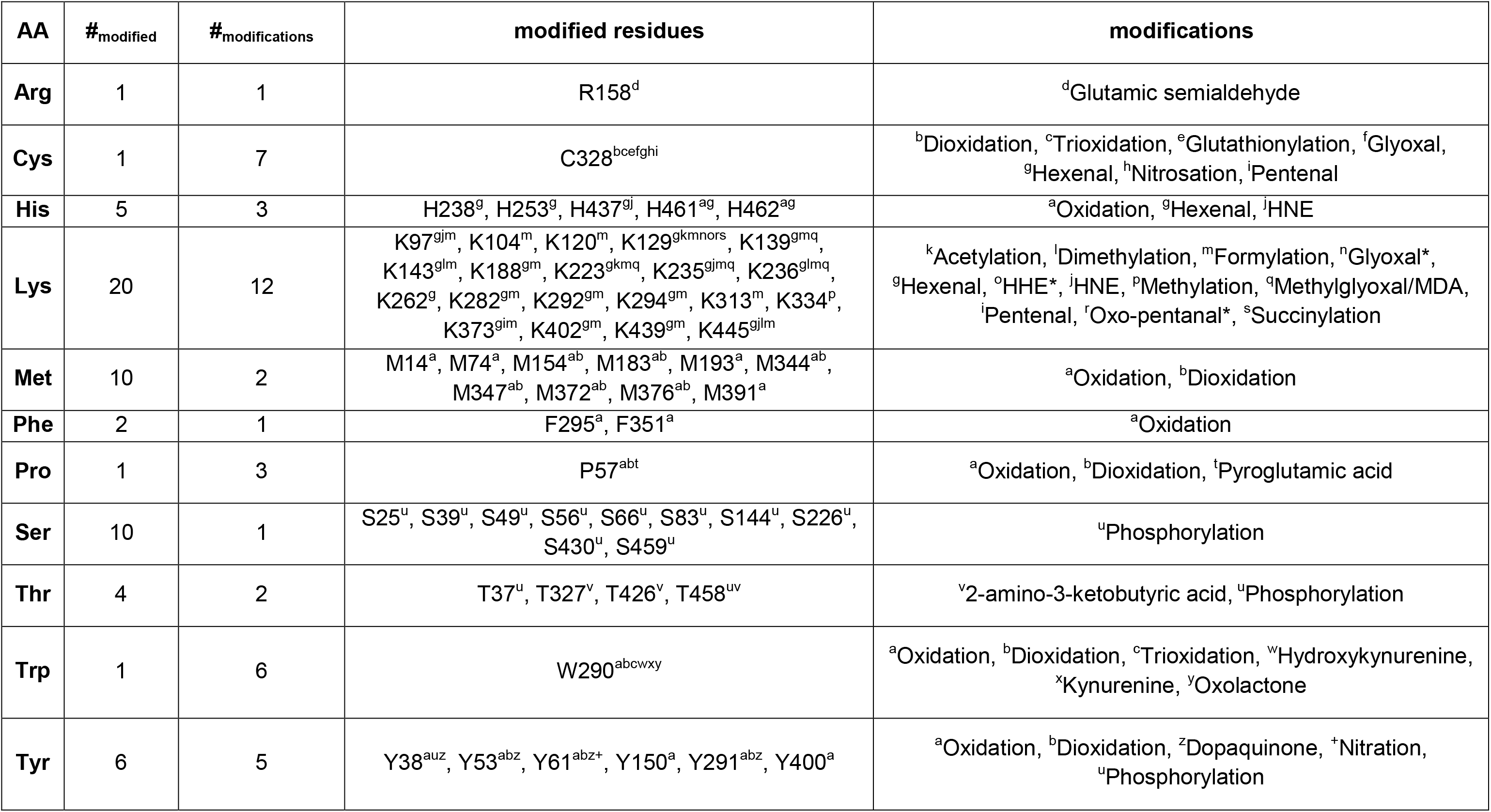
Vimentin modification sites identified in SIN-1 treated and untreated cardiac cells. * Indicates modifications via Schiff base formation.

**Figure 2:**
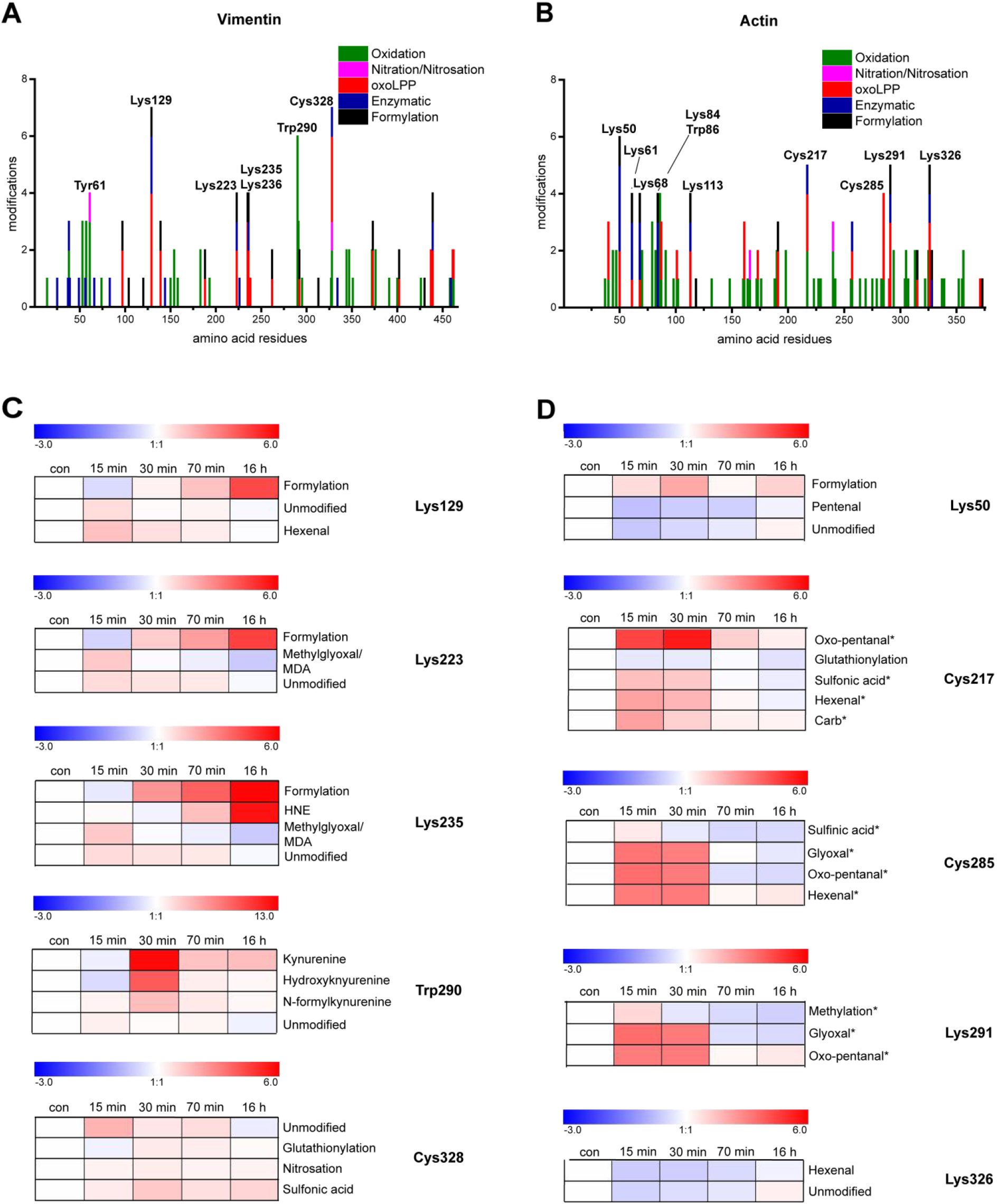
Summary of results obtained from PTM profiling and label-free relative quantification. Diagrams representing the number and types of modifications along the amino acid sequence of (A) vimentin and (B) actin. Heat maps illustrate the dynamics of different PTMs identified on (C) vimentin and (D) actin PTM “hot spots”; *indicates the presence of methionine sulfoxide on the peptide covering the modification site.

### Actin PTMs show modification specific dynamics

Modified peptides corresponding to six actin isoforms (cytoplasmic 1, cytoplasmic 2, alpha cardiac muscle 1, aortic smooth muscle, gamma-enteric smooth muscle and alpha skeletal muscle) were detected. Overall, 415 peptides carrying 40 different PTMs were identified (Table 2 and S4).

**Table 2:**
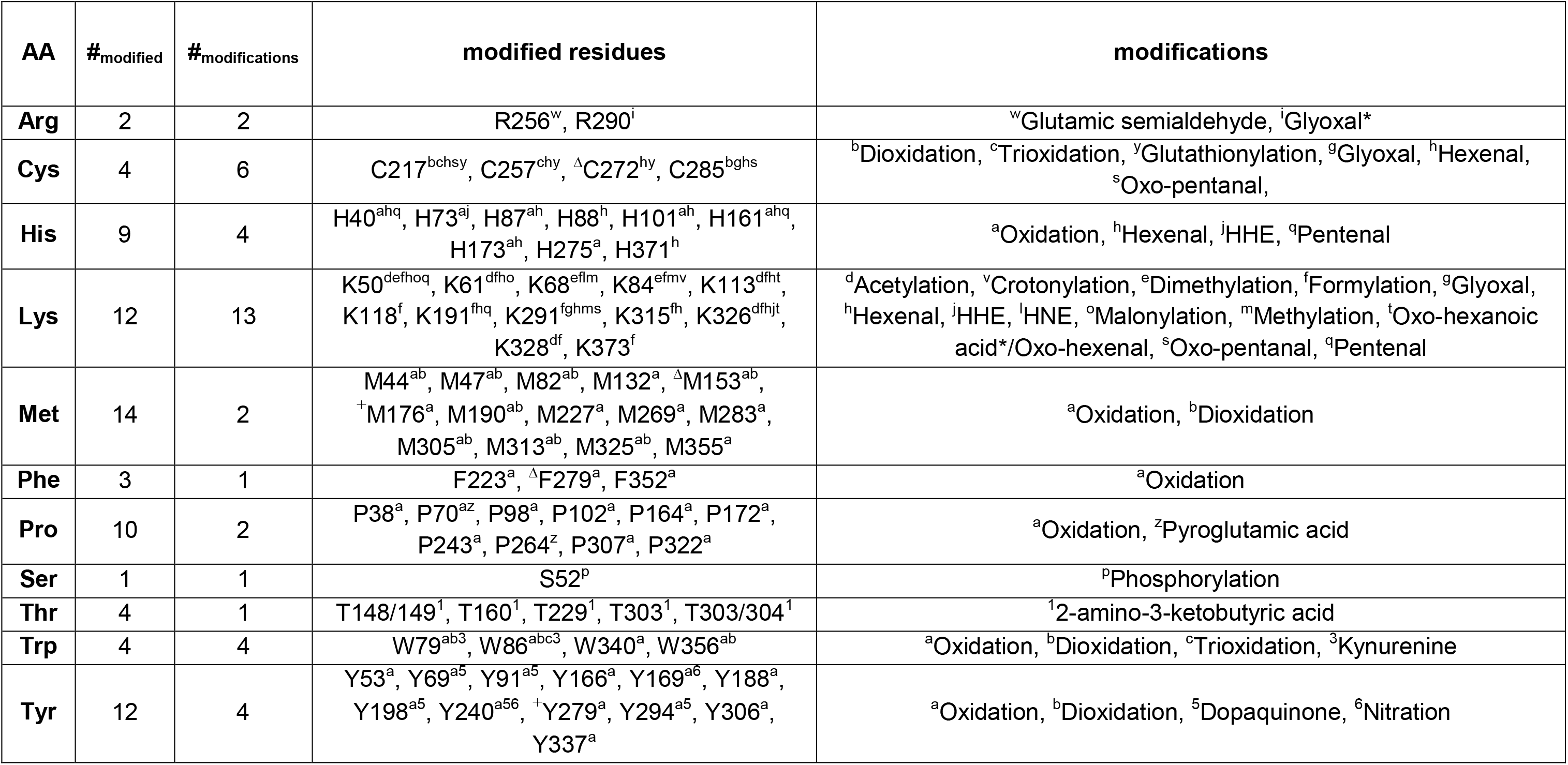
Actin modification sites identified in SIN-1 treated and untreated cardiac cells. Modification sites are numbered for the mature protein without propeptide as provided by UniProtKB. *indicates modifications via Schiff base formation. ^Δ^cytoplasmic actin isoforms: P60711, P63259. ^+^actin isoforms: P68035, P62738, P63269, P68136.

As observed in vimentin, a few residues represented highly populated PTM sites (Figure 2B). Interestingly, three Cys residues were glutathionylated (Cys217, 257 and 272). Of these, Cys217 emerges as a hot spot being detected in five modified states, including dioxidation and trioxidation, glutathionylation, and modification by hexenal and oxo-pentanal. Remarkably, glutathionylation of Cys217 was higher under basal conditions, and underwent a sustained decrease from the first time point after SIN-1 treatment (Figure 2D, Table S5). Cys285 was modified by four different moieties, also including dioxidation, hexenal and oxo-pentanal. Cys374 was not observed, likely due to the small size of the peptide.

Regarding Lys residues, Lys50 was detected in six forms including modifications by acetylation, dimethylation, malonylation, formylation, hexenal and pentenal adducts. Lys291 and Lys326 were found in five different modified forms, including methylation or acetylation, and formylation and modification by various reactive aldehydes. In addition, Lys61, Lys68 and Lys113 presented four modifications each. Again, several of the modifications affecting hot spot residues, including the modifications of Cys217, 285 and Lys291 by reactive aldehydes, showed a transient pattern, with higher levels at 15 and 30 min after SIN-1 addition, and returned to basal levels, or even below, at later times (Figure 2D).

Several Trp residues suffered various oxidative modifications and Trp86, in particular, presented four oxidative PTMs, including oxidation, dioxidation and trioxidation and formation of kynurenine.

Remarkably, the most abundant modification was Met oxidation to sulfoxide (14 residues) or sulfone (eight residues). Nevertheless, diverse oxidative and/or electrophilic modifications were found in numerous Lys residues (13 different PTMs distributed in 12 out of 19 positions), Tyr, Pro and His residues (see Table 2 for a detailed description). A single site of phosphorylation was detected on Ser52.

Hierarchical clustering analysis of actin modifications revealed several clusters of PTMs with similar dynamics (Figure S7B). Most of the direct oxidation sites as well as Lys formylations increased at early experimental time points and decreased after 16 h. That was also true for most of the Cys, Arg, and His oxoLPP adducts. However, hexenal and pentenal modifications on Lys and His residues showed a completely different behavior with a significant decrease in abundance after 15, 30 and 70 min of SIN-1 treatment. These PTMs also formed a distinct cluster visualized by PCA loading plots showing opposite projection in comparison to the majority of the oxidation sites (Figure S9). In contrast, Cys and Arg residues modified by oxoLPPs showed different dynamics with a maximum at 15 or 30 min. Remarkably, their corresponding tryptic peptides contained also methionine sulfoxide suggesting that the strong increase is a result of the oxidative state of the peptide. HNE-modified Lys68 (without Met sulfoxide) increased until 70 min of SIN-1 treatment and decreased below control levels after 16 h. Interestingly, the dynamics of hexenal- and HNE-adducts showed opposite behavior in vimentin and actin.

### Tubulins are apparently modified to a lesser extent than the other cytoskeletal structures in the nitroxidative stress model

Modified peptides corresponding to six α-tubulins (α-1A, α-1B, α-1C, α-3, α-4A, and α-8; Table S6), five β-tubulins (β-2A, β-2B, β-3, β-4B, and β-5; Table S7), and γ-tubulin (Table S8) were identified. Overall, ten modification sites were localized on α-tubulins, of which Met oxidation to sulfoxide (positions 154, 302, 377, 398, 413, and 425) was the most prominent one. Additionally, dioxidation of Met377 to sulfone, formylation of Lys 96, phosphorylation of Thr337, glutathionylation of Cys347, and nitrosation of Cys376 were detected (Table 3 and S6, Figure S3).

**Table 3:** **α,β and γ**-tubulin modification sites identified in SIN-1 treated and untreated cardiac cells.

| AA | #<sub>modified</sub> | #<sub>modifications</sub> | modified residues | modifications |
| --- | --- | --- | --- | --- |
| **α-tubulin** |  |  |  |  |
| **Cys** | 2 | 2 | C347<sup>c</sup>, C376<sup>d</sup> | <sup>c</sup>Glutathionylation, <sup>d</sup>Nitrosation |
| **Lys** | 1 | 1 | K96<sup>g</sup> | <sup>g</sup>Formylation |
| **Met** | 6 | 2 | M154<sup>a</sup>, M302<sup>a</sup>, M377<sup>ab</sup>, M398<sup>a</sup>, M413<sup>a</sup>, M425<sup>a</sup> | <sup>a</sup>Oxidation, <sup>b</sup>Dioxidation |
| **Thr** | 1 | 1 | T337<sup>i</sup> | <sup>i</sup>Phosphorylation |
| **β-tubulin** |  |  |  |  |
| **His** | 1 | 2 | H264<sup>ae</sup> | <sup>a</sup>Oxidation, <sup>e</sup>Hexenal |
| **Lys** | 2 | 2 | K103<sup>f</sup>, K252<sup>g</sup> | <sup>f</sup>Dimethylation, <sup>g</sup>Formylation |
| **Met** | 14 | 2 | M1<sup>a</sup>, M73<sup>ab</sup>, M147<sup>a</sup>, M164<sup>a</sup>, M170<sup>a</sup>, M233<sup>a</sup>, M257<sup>ab</sup>, M267<sup>ab</sup>,<br>M293<sup>a</sup>, M321<sup>ab</sup>, M323<sup>ab</sup>, M330<sup>a</sup>, M363<sup>a</sup>, M388<sup>a</sup> | <sup>a</sup>Oxidation, <sup>b</sup>Dioxidation |
| **Phe** | 2 | 1 | F265<sup>a</sup>, F389<sup>a</sup> | <sup>a</sup>Oxidation |
| **Pro** | 6 | 1 | P70<sup>a</sup>, P160<sup>a</sup>, P171<sup>a</sup>, P259<sup>a</sup>, P268<sup>a</sup>, P272<sup>a</sup> | <sup>a</sup>Oxidation |
| **Ser** | 1 | 1 | S322<sup>i</sup> | <sup>i</sup>Phosphorylation |
| **Thr** | 2 | 1 | T136<sup>j</sup>, T166<sup>j</sup> | <sup>j</sup>2-amino-3-ketobutyric acid |
| **Trp** | 2 | 4 | W101<sup>abkl</sup>, W344<sup>abk</sup> | <sup>a</sup>Oxidation, <sup>b</sup>Dioxidation, <sup>k</sup>Kynurenine,<br><sup>l</sup>Oxolactone |
| **Tyr** | 1 | 1 | Y200<sup>m</sup> | <sup>m</sup>Dopaquinone |
| **γ-tubulin** |  |  |  |  |
| **Met** | 1 | 1 | M377/8<sup>b</sup> | <sup>b</sup>Dioxidation |

β-tubulin showed a higher variety of PTMs. In total, 31 modification sites were identified including 14 Met, six Pro, two Lys, two Trp, two Thr, two Phe, one Tyr, one Ser and one His residues (Table 3 and S7, Figure S4). Most of the PTMs detected were of oxidative origin. Similar to α-tubulin, Met oxidation to sulfoxide was the most abundant PTM. Additionally, five Met residues (positions 73, 257, 267, 321, and 323) were further oxidized to sulfones. Hydroxyproline was the second most abundant modification occupying positions 70, 160, 171, 259, 268, and 272. Trp 101 and 344 were oxidized to hydroxy-Trp/oxindolylalanine, *N*-formylkynurenine/dioxindolylalanine, and kynurenine. Additionally, oxidation of Trp101 to oxolactone was detected. Phe265 and 389 were hydroxylated and Thr136 and 166 were present in carbonylated form (2-amino-3-ketobutyric acid). Tyr200 was transformed into DQ. His264 was oxidized and modified by hexenal. Lys modifications included dimethylation (Lys103) and formylation (Lys252). Only one phosphorylation site was detected on Ser322.

For γ-tubulin only one modification site, sulfone on Met377/378, was identified (Table 3 and S8).

For tubulins the only significantly SIN-1 regulated peptides belonged to α-tubulin and covered Cys347 and 376. Thus, peptides containing Cys347 in its glutathionylated form increased at 30 min upon stress induction and a peptide with S-nitrosated Cys376 was slightly elevated at 15, 70 min and 16 h (Table S9).

## Discussion

The cytoskeleton is a dynamic superstructure capable of responding to changes in the extra- and intracellular environments. Alterations in redox balance are involved in numerous physiological and pathological events (31,32). A scientific paradigm on the role of ROS/RNS in biological systems underwent significant renewal over the last decades. ROS/RNS, previously considered as deleterious by-products of metabolism or results of exogenous stressors (e.g. ionizing radiation), are currently acknowledged as essential messengers in signal transduction pathways and cell homeostasis (33).

A large variety of non-enzymatic PTMs are believed to be the driving force of redox signaling and lipoxidative stress regulated events (15,21,34,35). These PTMs include amino acid residue oxidation, protein carbonyl formation, nitration, nitrosation and addition of oxidized lipids (lipoxidation) and carbohydrate metabolites, and many of them can occur under basal and stress conditions. Moreover, these PTMs co-exist *in vivo* with all other types of enzymatic modifications including phosphorylation, methylation, acylation, ubiquitination and others. Furthermore, substantial cross-talk exists between different types of modification since many enzymes catalyzing PTMs are themselves targets for nitroxidative or lipoxidative stress and related PTMs. In order to understand multiple PTM cross-talk in biological systems, the dynamics of PTMs need to be studied. Thus, we designed and characterized a dynamic model of mild nitroxidative stress using rat primary cardiac cells treated with a peroxynitrite donor (SIN-1) for increasing times to monitor its effects on cytoskeletal dynamics and PTM cross-talk.

Vimentin, actin and tubulins are key players in cytoskeletal organization and dynamics. These proteins are major targets of non-enzymatic PTMs in physiological and pathophysiological conditions associated with redox imbalance (13,36,37). In our model, mild nitroxidative stress induced fast cytoskeletal reorganization. Vimentin filaments underwent a retraction from the cell periphery with formation of perinuclear condensed structures. A similar remodeling has been described in response to several types of stress, including exposure of endothelial cells to hypoxic conditions (38) and treatment of rat mesangial cells with several electrophilic molecules (39,40). Actin microfilaments switched from prominent stress fibers to cortical actin, with loss of cytoplasmic polymerized actin. This is in accordance with previous works reporting a decrease in F-actin in response to peroxynitrite (41). In turn, microtubules changed from homogenously distributed filaments to juxtanuclear condensed structures and thinner peripheral staining, similar to the effects observed in connection with treatment with cyclopentenone prostaglandins (39), changes in mitochondrial dynamics (42), and autophagic degradation of aggregated proteins (43).

Importantly, cytoskeletal remodeling appeared to be transient and 16 h after stress induction cytoskeletal networks recovered their normal morphology.

Overall, effects of nitroxidative stress on cytoskeletal remodeling can occur directly through PTMs or indirectly through disrupted interaction with binding partners and (de-)activation of signaling pathways regulating cytoskeletal proteins. Using dedicated enrichment and separation techniques combined with high-resolution mass spectrometry we performed in-depth proteomics profiling of vimentin, actin and tubulin PTMs covering over 60 different PTM types including enzymatic, oxidative and oxoLPP-based modifications. Cytoskeletal proteins showed different susceptibility to PTMs. Curiously, relatively few modification sites and types were detected for tubulins, whereas vimentin and actin were heavily modified.

Surprisingly, Lys formylation was the most abundant modification in actin and vimentin, whereas only two formylated residues were found in α- and β-tubulins. Furthermore, formylation in actin and vimentin had different dynamics. While vimentin formylation was maximal after 16 h of SIN-1 treatment, the highest formylation of actin occurred between 15 and 30 min. Formylation has been previously reported to occur during *in vitro* incubation with peroxynitrite (44), which might explain the high number of identified modification sites. However, the differences in formylation abundance and dynamics observed among the different proteins studied indicate that it is not a chemical artefact, randomly modifying all Lys-residues, but rather a naturally occurring modification. Indeed, previous reports of Lys-formylation include the modification of nuclear proteins via reaction with formyl-phosphate, generated by oxidative DNA damage (45,46) and the formation of N(6)-formyl lysine by exposure to endogenous formaldehyde (47). Furthermore, formyl-CoA, produced in peroxisomes via α-oxidation of methyl-substituted fatty acids (48), is also considered as an intracellular donor of the formyl group (49). Although the functional role of Lys formylation is not completely understood, it is considered that it could interfere with acetylation or other Lys modification. Moreover, as other modifications altering the charge of Lys residues, it could interfere with protein-protein interactions (50). Interestingly, enzymes capable of reversing formylation have been identified (51), which together with the high abundance and dynamic nature of this PTM make it an interesting target.

Hexenal, a product of polyunsaturated fatty acid oxidation, which accumulates in this model (28), was another abundant moiety targeting most of the nucleophilic residues in actin and vimentin. Hexenal-protein adducts had been previously demonstrated both in *in vit*ro co-incubation experiments (52,53), as well as in biological samples (54). Nevertheless, it is important to note that assignment of hexenal adducts in this study was based on the identification of a certain mass increment corresponding to the elemental composition C6H12O and does not provide structural confirmation. Interestingly, there were clear differences in the dynamics of oxoLPP-adducts between Lys, His, Cys, and Arg residues. For instance in actin, hexenal-adducts on Lys 50, 61, 315, 326 and His 40 residues decreased between 15 and 70 min of SIN-1 treatment, whereas hexenal-adducts of Cys217 and 285 significantly increased after 15 min. However, it should be considered that peptides with oxoLPP modified Cys residues also contained Met sulfoxide, which might contribute to the observed dynamics. Indeed, peptides covering hexenal modified Lys191 or His161 also contained Met sulfoxide and showed dynamics similar to Cys containing peptides. Moreover, adducts of different oxoLPPs demonstrated different behaviour, e.g. hexenal- and pentenal-adducts versus HNE, HHE and methylglyoxal/MDA. Furthermore, dynamics for most of the PTMs were protein-specific, indicating its importance in the regulation of cytoskeletal dynamics upon nitroxidative stress.

An important implication of our observations is the transient nature of both, the morphological effects observed, as well as of many of the PTMs identified. Several PTMs are readily and rapidly reversible. However, in other cases, for instance lipoxidation, the stability of the adducts will depend on many factors, including their chemical environment and further reactivity (55), as well as the presence of enzymatic mechanisms capable of catalyzing the reverse reaction, such as thioredoxins or sirtuins (56,57). In addition, some modifications can be considered irreversible. In those cases, the reversibility of the effect would probably rely on protein turnover. Importantly, the bottom-up proteomic approach employed here provides unprecedented detailed information on the PTM landscape; however, it does not provide information on the variety of proteoforms arising under these conditions, that is, how PTMs are clustering on single protein molecules in order to generate diverse functional species, or whether the decay of specific PTMs is related to reversibility or protein turnover. Therefore, understanding the intimate mechanisms underlying the transient effects observed and the underlying PTM crosstalk will require additional efforts.

Several amino acid residues in actin and vimentin, including Lys50 and Cys217 in actin and Lys129 and Cys328 in vimentin, showed higher susceptibility to a wide variety of PTMs, representing PTM hot spots, which might play an important role in direct PTM cross talk. In contrast, fewer residues in tubulins presented several modifications, of which Trp 101 and 344 displayed the highest diversity with four and three modifications, respectively (Table 3).

Several of the hot spots identified for vimentin and actin were Lys residues. Curiously, Lys was not detected in its oxidized forms (aminoadipic semialdehyde or aminoadipic acid), possibly due to their previous/preferential modification by other moieties. Besides the variety of oxoLPP adducts we also identified enzymatic Lys PTMs, including acetylation, mono- and dimethylation, malonylation, succinylation, and crotonylation. Unfortunately, acetylation of Lys 40, reported to occur under various pathophysiological conditions, could not be assessed due to the large size of the corresponding tryptic peptide.

Besides Lys, Cys-, Trp-, and Tyr-residues were modified by multiple PTMs as well. Reversible Cys modifications including nitrosation, glutathionylation and oxidation to sulfenic acid or disulfides, are considered redox switches. These modifications reversibly regulate protein function and are at the basis of redox signaling (58,59). Notably, other thiol modifications, including irreversible oxidations and addition of oxoLPPs, can occur on Cys residues leading to disruption or modulation of redox signaling (17,21). Importantly, the structural diversity of Cys modifications contributes to the generation of a wide variety of protein functional species. Identified Cys hot spots in vimentin and actin displayed various oxidative and oxoLPP modifications, which showed different dynamics on the same residue. Remarkably, lipoxidation of cysteine residues in vimentin and actin has been previously reported in several pathophysiological models (60–63) (see (13,18) for review).

The identified Tyr hot spot in vimentin, Tyr61, was modified by oxidation, dioxidation, nitration and transformation into DQ. Taking into account the current indication of a possible reversibility of Tyr nitration (64,65), this modification site might represent an interesting case for further evaluation.

It is also interesting that, six out of ten actin PTM hot spots lied within a 53 amino acid sequence, indicating not only possible cross-talk by direct competition of different PTMs for the same residue, but also influence of one modification site to the others. Indeed, covalent linkage of large and often charged functional groups might induce significant changes in the molecular environment, thus forcing the surrounding atoms to optimize their interactions, a phenomenon known as “population shift” (66). Nevertheless, top-down proteomic approaches are needed to obtain information on the distribution of modified species.

From a functional point of view, few of the hot spots identified here have been the subject of targeted studies. Cys328 of vimentin is critical for remodeling of the network in response to oxidants and electrophilic mediators, as evidenced from the blunted or attenuated reorganization of cysteine-deficient mutants (40,67). Moreover, the presence of vimentin cysteine is key for normal filament assembly and dynamics since a Cys328Ser vimentin mutant forms filaments with significantly different kinetics and morphology, and increased resistance to disruption even by agents that are not exclusively cysteine-specific, suggesting that cysteine modification may be required for reorganization in response to different types of stress (67,68). Therefore, the variety of the modifications detected on Cys328 and their dynamic nature strengthen our previous observations on its role as a sensor of oxidative and electrophilic stress (40,67).

Interestingly, our results show a temporal correlation between cytoskeletal remodeling and some of the PTMs identified, providing a basis to address their functional significance. To this point, PTM hot spots identified here in vimentin and actin represent first targets for site-directed mutagenesis to understand their role in PTMs cross-talk and redox regulation.

## Abbreviations

CID: collision-induced dissociation
DDA: data dependent acquisition
DOPA: dihydroxyphenylalanine
DQ: dopaquinone
ESI: electrospray ionization
ETD: electron-transfer dissociation
HHE: hydroxyl-hexenal
HNE: hydroxy-nonenal
LPP: lipid peroxidation products
MDA: malondialdehyde
PCA: principal component analysis
PTM: posttranslational modification
RNS: reactive nitrogen species
ROS: reactive oxygen species
SIN-1: 3-morpholinosydnonimine
TMT: tandem mass tag

## Author Contributions

MF and DPS contributed to the idea conception and design of the experiments. EG performed all experiments, including data analysis, except Cys TMT labeling. VV performed Cys TMT labeling experiments. EG and AM performed cell microscopy acquisition and image analysis. The manuscript was written through the contributions of all authors. All authors have given approval to the final version of the manuscript.

## Acknowledgements

Feedback from EU COST Actions EuroCellNet (CA15214) and EpiLipidNet (CA19105) is gratefully acknowledged.

