## Supplementary data for "Dynamic posttranslational modifications of cytoskeletal proteins unveil hot spots under nitroxidative stress"

**Running Title:** PTMs hot spots in cytoskeletal proteins

[a]Present address: Drug Discovery Sciences, Boehringer Ingelheim Pharma GmbH & Co. KG, Biberach an der Riss, Germany

[b]Present address: Linnaeus University, Dept. of Chemistry and Biomedical Sciences, Universitetskajen, 391 82 Kalmar, Sweden

**Table of contents**

**Supplementary materials and methods** ........................................................................ **S3-8**

**Figure S1**: Gel electrophoresis prior LC-MS/MS analysis. ................................................. **S9**

**Figure S2**: Schematic representation of LC-MS/MS analysis strategy for identification of modified peptides in actin, vimentin and tubulin. ................................................................ **S10**

**Figure S3:** Sequence alignment of identified α-tubulin isoforms with modified residues labeled. ............................................................................................................................... **S11**

**Figure S4:** Sequence alignment of identified β-tubulin isoforms with modified residues labeled. ............................................................................................................................... **S12**

**Figure S5:** Amino acid sequence of vimentin with modified residues labeled. .................. **S13**

**Figure S6:** Sequence alignment of identified actin isoforms with modified residues labeled. ............................................................................................................................... **S14**

**Figure S7**: Hierarchical clustering of modification sites in vimentin and actin based on the relatively quantified corresponding tryptic peptides. ......................................................... **S15**

**Figure S8**: PCA loadings plot of modification sites in vimentin based on the relatively quantified corresponding tryptic peptides. ........................................................................ **S16**

**Figure S9**: PCA loadings plot of modification sites in actin based on the relatively quantified corresponding tryptic peptides. ......................................................................................... **S17**

#### Supplementary materials and methods

**Materials** – Primary rabbit polyclonal anti-actin (20-33) antibody (A5060), ammonium hydroxide (25% w/v), lactic acid (85%-90% w/v), pyrrolidine, trifluoroacetic acid (TFA), neocuproine, methyl methanethiosulfonate (MMTS), reduced glutathione, glutathione reductase (GR), and all ammonium and sodium salts were purchased from Sigma-Aldrich GmbH (Taufkirchen, Germany). SDS, urea, methanol and dithiothreitol (DTT) were obtained from Carl Roth GmbH + Co. KG (Karlsruhe, Germany). HEPES was purchased from Merck Chemicals GmbH, Darmstadt, Germany, Tris, iodoacetamide, and EDTA were from Applichem GmbH (Darmstadt, Germany) and phosphatase-inhibitor-mix solution and CHAPS were purchased from Serva Electrophoresis GmbH (Heidelberg, Germany). Glutaredoxin 1 from E. coli (mutant C14S) was bought from IMCO Corporation Ltd AB, (Stockholm, Sweden), NADPH, tetrasodium salt was purchased from Biomol GmbH (Hamburg, Germany) and sodium L-ascorbate was from Santa Cruz Biotechnology, Inc., (Heidelberg, Germany). Pierce $TiO_2$ phosphopeptide enrichment and clean-up kit, iodoTMTsixplex isobaric label reagent set, immobilized anti-TMT antibody resin, TMT elution buffer and Zeba Spin desalting columns were from Thermo Scientific (Life Technologies GmbH, Darmstadt, Germany) and peroxidase-conjugated goat anti-rabbit antibody from Jackson ImmunoResearch Laboratories, Inc. (West Grove, PA, USA). Protein free blocking solution (AdvanBlock), washing solution (AdvanWash) and WesternBright Sirius HRP substrate were obtained from Advansta Inc. (Biozym Scientific GmbH, Hessisch Oldendorf, Germany).

**Western Blot analysis of actin** - Proteins in lysis buffer (10 µg) were mixed with Laemmli sample buffer (62.5 mmol/L Tris-HCl pH 6.8, 20% v/v glycerol, 2% w/v SDS, 5% v/v ß-mercaptoethanol, 0.01% w/v bromophenol blue) and separated by SDS-PAGE (12% T, 0.75 mm, 200 V; BioRad mini protean III cell; BioRad Laboratories GmbH, München, Germany). Proteins were tank blotted onto a PVDF membrane in Bjerrum Schafer-Nielsen transfer buffer (48 mmol/L Tris, 39 mmol/L glycine) using Mini Trans-Blot Cell (60 min, 100 V; BioRad). Membranes were blocked 1 h (RT, protein free blocking solution), incubated with primary rabbit polyclonal anti-actin (20-33) antibody (1 h, 1:10,000 in blocking solution, RT) and washed (10 min, three times, washing solution). Afterwards membranes were incubated with peroxidase-conjugated goat anti-rabbit antibody (1 h, 1:10,000 in blocking solution, RT) followed by washes with washing solution (10 min, two times) and Tris-buffered saline (TBS, 20 mmol/L Tris, 500 mmol/L NaCl; 10 min). Membranes were developed with enhanced chemiluminescence (ECL) using WesternBright Sirius HRP substrate. Images were taken with the Fusion FX7 imaging system (Peqlab Biotechnologie GmbH, VWR International GmbH, Erlangen, Germany).

**Protein extraction for phophoproteomics** - When cells reached 80% confluence they were treated with 10 µmol/L SIN-1 for 15 min, 30 min, 70 min and 16 h in serum-free medium (DMEM/F12 supplemented with 100 U/mL penicillin and 100 µg/mL streptomycin). After treatment cells were washed with warm PBS and harvested by trypsinization followed by centrifugation (4 min, 300 × *g*, 4 °C). Cell pellets were washed with ice cold PBS (two times, 10 min, 1000 × *g*, 4 °C) and snap-frozen at -80 °C. Cell pellets were resuspended in lysis buffer (7 mol/L urea, 2 mol/L thiourea, phosphatase-inhibitor-mix (1:100) in 50 mmol/L Tris-HCl, pH 7.4). Samples were sonicated using a Vibra-Cell tip sonicator (20 kHz, 1 min with on/off pulse 5 s each, 40% amplitude; Sonics & Materials, Inc., Newtown, CT, USA) followed by centrifugation (20 min, 10,000 × *g*, 4 °C). Supernatants were collected and protein concentration was determined by Bradford assay (1).

**In-solution tryptic digestion for phosphoproteomics** - Proteins (800 µg) were reduced with 5 mmol/L TCEP (90 min, 37 °C, 550 rpm), alkylated with 10 mmol/L IAA (60 min, 27 °C, 550 rpm, in the dark) and excess of IAA was quenched with 10 mmol/L DTT (30 min, 37 °C, 550 rpm). Samples were diluted with 50 mmol/L $NH_4HCO_3$ to a final concentration of 1 mol/L urea and digested with trypsin overnight (1:50 trypsin to protein ratio, in 3 mmol/L $NH_4HCO_3$, 37 °C, 550 rpm). Afterwards samples were desalted with solid-phase extraction using Waters Oasis HLB 1cc (10 mg). The stationary phase was rinsed with methanol (1 mL) and equilibrated with water (1 mL). Samples were loaded, stationary phase was washed with 0.1% formic acid in 7% aqueous acetonitrile (3x, 1 mL) and peptides were eluted with 0.5% formic acid in 70% aqueous acetonitrile (500 µL). Eluates were vacuum concentrated.

**Enrichment of phosphopeptides with $TiO_2$** - Enrichment was performed using Pierce $TiO_2$ phosphopeptide enrichment and clean-up kit according to the manufacturer's protocol with slight changes. $TiO_2$ spin tips were washed with solution A (80% acetonitrile, 0.1% TFA; 20 µL, 3000 × *g*, 2 min) and solution B (57% v/v acetonitrile, 28.6% v/v lactic acid, 0.1% TFA, 20 µL). Phosphopeptides were resuspended in 150 µL solution B and applied to $TiO_2$ spin tips (1000 × *g*, 10 min). Flow-through of the sample was reapplied followed by washes with solution B (20 µL, 3000 × *g*, 2 min) and solution A (three times, 20 µL). Phosphopeptides were eluted with elution solution 1 (1.25% w/v $NH_4OH$) and elution solution 2 (5% v/v pyrrolidine; each 50 µL, 1000 × *g*, 5 min). Eluted phosphopeptides were brought to pH 2.0-2.5 with 150 µL 2.5% TFA.

**Graphite clean-up of enriched phosphopeptides** – Storage buffer of graphite spin columns was removed by centrifugation (2000 × *g*, 1 min). Graphite columns were activated by 1 mol/L $NH_4OH$ (two times), 100% acetonitrile and 1% TFA (two times, 100 µL each). Phosphopeptides were added and periodic vortex mixing (10 min) allowed their binding. Columns were centrifuged (1000 × *g*, 3 min) and washed with 1% TFA (200 µL, 2000 × *g*, 1 min, two times). Phosphopeptides were eluted with 0.1% v/v formic acid in 50% v/v acetonitrile (100 µL, 2000 × *g*, 1 min, four times). Combined eluates were dried by vacuum concentration and stored at -20 °C. Before MS analysis peptides were dissolved in 40 µL of 0.1% formic acid in 3% aqueous acetonitrile.

**Mass Spectrometry of phosphopeptides** - A nano-Acquity UPLC (Waters GmbH, Eschborn, Germany) was coupled online to an LTQ Orbitrap XL ETD mass spectrometer equipped with a nano-ESI source (Thermo Fischer Scientific, Bremen, Germany). Eluent A was aqueous formic acid (0.1% v/v), and eluent B was formic acid (0.1% v/v) in acetonitrile. Samples (10 µL) were loaded onto the trap column (nanoAcquity symmetry C18, internal diameter 180 µm, length 20 mm, particle diameter 5 µm) at a flow rate of 10 µL/min. Peptides were separated on BEH 130 column (C18-phase, internal diameter 75 µm, length 100 mm, particle diameter 1.7 µm) with a flow rate of 0.4 µL/min. Enriched peptides were separated using two step gradients from 3 to 35% eluent B over 180 min and then to 85% eluent B over 40 min. After 5 min at 85% eluent B the column was equilibrated for 15 min and samples were injected every 240 min. The transfer capillary temperature was set to 200 °C and the tube lens voltage to 120 V. An ion spray voltage of 1.5 kV was applied to a PicoTip online nano-ESI emitter (New Objective, Berlin, Germany). The precursor ion survey scans were acquired at an orbitrap (resolution of 60,000) for a *m/z* range from 400 to 2000. ETD-tandem mass spectra (irradiation time 100 ms, isolation width 2 u) were recorded in the linear ion trap by data-dependent acquisition (DDA) for the top six most abundant ions in each survey scan with dynamic exclusion for 60 s using Xcalibur software (version 2.0.7).

**Database search for phosphoproteins** - The acquired ETD tandem mass spectra were searched against the Uniprot *Rattus Norvegicus* database (downloaded on November, 17 2016, entries: 7,969 proteins) using Sequest search engine (Proteome Discoverer 1.4, Thermo Scientific), allowing up to two missed cleavages and a mass tolerance of 10 ppm for precursor ions and 0.8 Da for product ions. Oxidation of Met, Cys and Trp, carbamidomethylation of

Cys and phosphorylation of Ser, Thr and Tyr were used as variable modifications. Only peptides with high confidence, charge-dependent scores (Xcorr ≥2.0, 2.25, 2.5 and 2.75 for charge states 2, 3, 4 and 5) and ranked on position 1 were considered. False discovery rates (FDR) were set below 1%. Additionally phosphorylation site probabilities were determined using PhosphoRS 3.0 (within Proteome Discoverer 1.4) (2). Only phosphopeptides with PhosphoRS scores ≥50 and phosphorylation sites with probabilities ≥75% were considered. For ambiguous localization sites all possible residues were included.

**Protein extraction for iodoTMTsixplex labeling of reversibly modified cysteines -** After SIN-1 treatment, cells were washed, scraped into ice cold PBS, and collected by centrifugation (10 min, 1000 × *g*, 4 °C). Cell pellets were washed again with ice cold PBS (two times) and resuspended in HENS buffer (1 mmol/L EDTA, 0.1 mmol/L neocuproine, 2% SDS, 50 mmol/L methyl methanethiosulfonate (MMTS) in 150 mmol/L HEPES, pH 7.3). Samples were sonicated on ice using a Vibra-Cell tip sonicator (20 kHz, 2 min with on/off pulse 5 s each, 40% amplitude; Sonics & Materials, Inc., Newtown, CT, USA) and centrifuged (20 min, 10,000 × *g*, 4 °C). Supernatants were collected and reduced cysteines (free thiols, -SH) were blocked with 50 mmol/L MMTS in HENS buffer (added fresh; 30 min, 37 °C, 350 rpm) followed by centrifugation (10 min, 20,000 × *g*, 4 °C). Excess of MMTS was removed by protein precipitation using ice-cold acetone (five volumes, 1 h, -20 °C) followed by centrifugation (30 min, 10,000 × *g*, RT), washes with 1 mL of ice-cold acetone, and vacuum concentration. Protein pellets were resuspended in AENS buffer (50 mmol/L $NH_4HCO_3$, 1 mmol/L EDTA, 0.1 mmol/L neocuproine, 2% CHAPS). Protein concentrations were determined by Bradford assay.

**IodoTMTsixplex labeling -** 1 mg/mL of proteins per condition was used for iodoTMT labeling. The whole procedure was carried out in the dark. S-glutathionylation was reduced enzymatically with 1 mmol/L glutathione, 1 mmol/L NADPH, 2.5 µg/mL glutaredoxin, 4 U/mL glutathione reductase (15 min, 37 °C, 500 rpm). Zeba Spin desalting columns were used to remove excess of reagents according to the manufacturer's instructions. IodoTMTsixplex reagents were prepared according to the manufacturer's instructions and added to samples. Specific reduction of S-nitrosation with 10 mmol/L sodium L-ascorbate and alkylation with iodoTMT reagents was carried out simultaneously. Five different protein samples were separately labeled using five different iodoTMTsixplex reagents (2 h, 37 °C, 500 rpm, in the dark). After iodoTMT labeling, equal volumes of each condition were pooled

(5 mg in 5 mL) and 10 volumes of ice cold acetone were added for protein precipitation at -20 °C for 2 h.

**In-solution tryptic digestion of iodoTMT labeled samples -** Protein pellets were resuspended in digestion buffer (50 mM $NH_4HCO_3$, 8 mol/L urea), reduced with 50 mmol/L DTT (90 min, 37 °C, 350 rpm) and free thiols were alkylated with 15 mmol/L IAA (60 min, 27 °C, 350 rpm, in the dark). Samples were diluted with 50 mmol/L $NH_4HCO_3$ to a final concentration of 1 mol/ L urea and digested with trypsin overnight (1:50 trypsin to protein ratio, in 3 mmol/L $NH_4HCO_3$, at 37 °C, 350 rpm). Samples were desalted by solid-phase extraction using Waters Oasis HLB 1cc (30 mg). Briefly, the stationary phase was rinsed with methanol (1 mL) and equilibrated with water (1 mL). Samples were loaded, washed (0.1% formic acid in 7% aqueous acetonitrile; 1 mL, three times) and peptides were eluted with 0.5% formic acid in 70% aqueous acetonitrile (500 µL) followed by vacuum concentration.

**Enrichment of iodoTMT-labeled peptides -** Approximately 5 mg iodoTMTsixplex labeled peptide mixture resuspended in Tris-buffered saline (TBS; 20 mmol/L Tris, 500 mmol/L NaCl) was enriched using anti-TM resin. The amount of anti-TMT resin required to enrich each replicate was estimated using the formula: resin slurry [µL] = 142.858 x sample amount (mg) (3). Peptide sample was incubated with anti-TMT resin on a rotary shaker for 2 h at RT. The unbound peptides were washed away by a series of wash steps (10 min each) including 2 mol/L urea in TBS (3 column volumes), 0.1% sodium deoxycholate (SDC) in TBS (3 column volumes), TBS (3 column volumes) and $H_2O$ (5 times, 4 column volumes each). Finally, iodoTMT labeled peptides were eluted from the resin with TMT elution buffer (4 column volumes). The eluates were acidified by adding 0.5% TFA and any residual precipitated SDC was removed by centrifugation (5 min, 20 000 × *g*, RT). IodoTMT labeled peptides were vacuum dried and stored at -20 °C. Desalting was performed by solid-phase extraction using Waters Oasis HLB 1cc (30 mg) as described before. Shortly before MS analysis the samples were dissolved in 25 µL of 0.1% formic acid in 3% aqueous acetonitrile.

**LC-MS/MS of iodoTMT labeled peptides -** A nano-Acquity UPLC system (Waters GmbH, Eschborn, Germany) coupled online to a Q-TOF Synapt G2-S*i* mass spectrometer, equipped with a nano-ESI source (Waters, MS Technologies, Manchester, UK) was used to analyse iodoTMT labeled tryptic peptides. Eluent A was aqueous formic acid (0.1% v/v), and eluent B was formic acid (0.1% v/v) in acetonitrile. 10 µL of each sample was loaded onto the trap column (nano-Acquity symmetry C18, internal diameter 180 µm, length 20 mm, particle diameter 5 µm) at a flow rate of 10 µL/min. Peptides were separated on a BEH 130 column

(C18-phase, internal diameter 75 µm, length 100 mm, particle diameter 1.7 µm, column temperature 30 °C) with a flow rate of 0.35 µL/min using a linear gradient from 3 to 30% eluent B in 89.5 min, and to 85% eluent B in 10 min. The doubly protonated signal of GluFib at *m/z* 785.8426 was acquired as lock mass. The precursor ion survey scan was acquired in positive ion mode with a resolution of 20,000 for an *m/z* range from 360 to 2000 using the following settings: capillary voltage 3.5 kV, source temperature 80 °C, sampling cone of 30 V, source offset of 60 V, cone gas flow of 20 L/h, purge gas flow of 600 mL/h, nanoflow gas pressure of 0.2 bar, and a scan time of 0.5 s. Collision-induced dissociation (CID) fragmentation occurred in the trap cell using a data-dependent acquisition (DDA) over the top 10 most intense signals in iTRAQ mode (elevated collision energy 30 V) with collision energy ramp from 50 to 70 V for *m/z* 1180. Tandem mass spectra were acquired in the *m/z* range of 50 to 2000 using MS/MS scan of 0.2 sec with a dynamic exclusion of ± 500 ppm within 30 sec. Data files (.RAW files) were processed with Progenesis QI for Proteomics 3.0 (version 2.1; NonLinear Dynamics, Newcastle upon Tyne, U.K.) to convert into ".mgf " format, using default settings.

**Database search of iodoTMT labeled peptides -** The acquired tandem mass spectra were searched against the Uniprot *Rattus norvegicus* database using Sequest search engine (Proteome Discoverer 1.4, Thermo Scientific), allowing up to two missed cleavages and a mass tolerance of 10 ppm for precursor ions and 0.8 Da for product ions. A list of variable modifications used for database search included oxidation, carbamidomethylation, and iodoTMTsixplex on Cys. The search filters used were rank 1 and high confidence. For the final dataset, peptides identified by MS/MS in at least two biological replicates were considered. The iodoTMTsixplex quantification method provided within Proteome Discoverer was used for quantification. The TMT ratios of reporter ions were quantified from MS2 scans using an integration tolerance of 20 ppm with the most confident centroid setting. Isotopic impurity factors provided by the manufacturer were used for correction of the reporter ion ratios.

#### Supplementary data

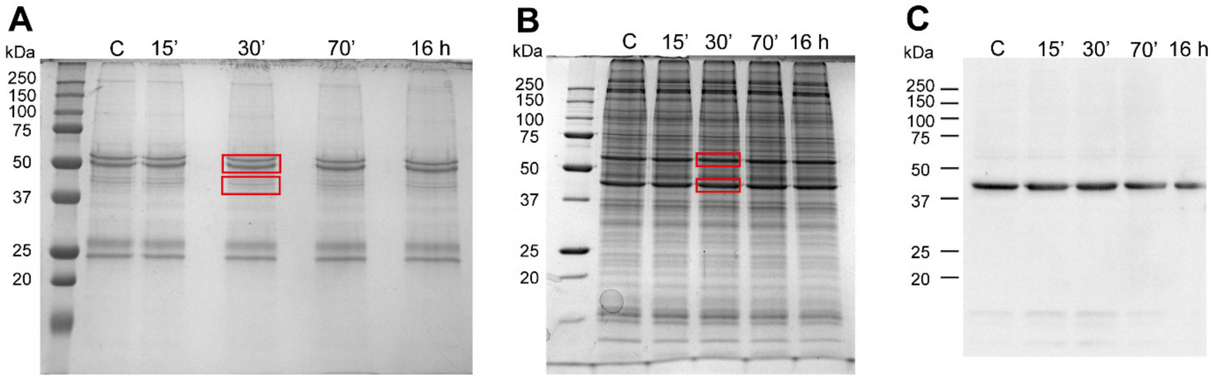


**Figure S1**: Gel electrophoresis prior LC-MS/MS analysis. (A) Immunoprecipitation of vimentin was performed using agarose conjugated anti-vimentin antibody. Eluted proteins were separated by SDS-PAGE. Two bands as labeled, containing vimentin as confirmed by MS analysis, were cut and in-gel-digested for subsequent LC-MS/MS analysis. (B) Proteins from cell extracts were separated by SDS-PAGE and bands corresponding to actin (42 kDa) and tubulin (55 kDa) were cut. (C) Western Blot analysis of actin to confirm localization of actin band on the corresponding SDS PAGE.

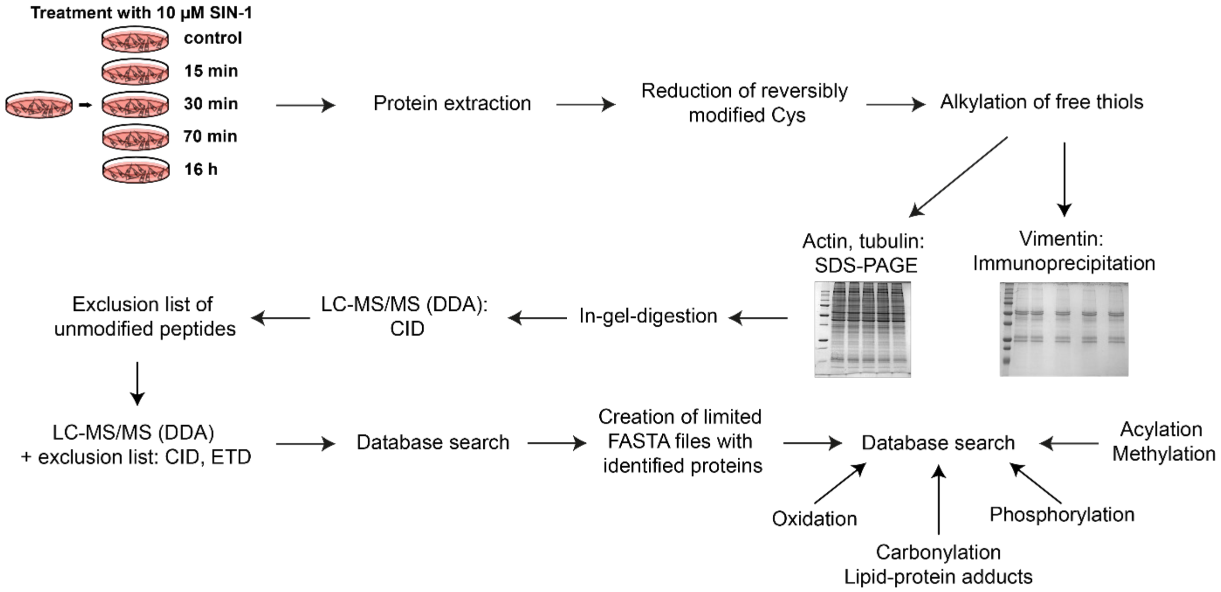


**Figure S2**: Schematic representation of LC-MS/MS analysis strategy for identification of modified peptides in actin, vimentin and tubulin.

#### α-TUBULIN

```
SP|P68370|TBA1A_RAT MRECISIHVGQAGVQIGNACWELYCLEHGIQPDGQMPSDKTIGGGDDSFNTFFSETGAGK 60
SP|Q6P9V9|TBA1B_RAT MRECISIHVGQAGVQIGNACWELYCLEHGIQPDGQMPSDKTIGGGDDSFNTFFSETGAGK 60
SP|Q6AYZ1|TBA1C_RAT MRECISIHVGQAGVQIGNACWELYCLEHGIQPDGQMPSDKTIGGGDDSFNTFFSETGAGK 60
SP|Q68FR8|TBA3_RAT  MRECISIHVGQAGVQIGNACWELYCLEHGIQPDGQMPSDKTIGGGDDSFNTFFSETGAGK 60
SP|Q5XIF6|TBA4A_RAT MRECISVHVGQAGVQMGNACWELYCLEHGIQPDGQMPSDKTIGGGDDSFTTFFCETGAGK 60
SP|Q6AY56|TBA8_RAT  MRECISVHVGQAGVQIGNACWELFCLEHGIQADGTFGTQASKINDDDSFTTFFSETGNGK 60
                    ******:********:*******:******* ** : :: :  ..****.***.*** **

SP|P68370|TBA1A_RAT HVPRAVFVDLEPTVIDEVRTGTYRQLFHPEQLITGKEDAANNYARGHYTIGKEIIDLVLD 120
SP|Q6P9V9|TBA1B_RAT HVPRAVFVDLEPTVIDEVRTGTYRQLFHPEQLITGKEDAANNYARGHYTIGKEIIDLVLD 120
SP|Q6AYZ1|TBA1C_RAT HVPRAVFVDLEPTVIDEVRTGTYRQLFHPEQLITGKEDAANNYARGHYTIGKEIIDLVLD 120
SP|Q68FR8|TBA3_RAT  HVPRAVFVDLEPTVVDEVRTGTYRQLFHPEQLITGKEDAANNYARGHYTIGKEIVDLVLD 120
SP|Q5XIF6|TBA4A_RAT HVPRAVFVDLEPTVIDEIRNGPYRQLFHPEQLITGKEDAANNYARGHYTIGKEIIDPVLD 120
SP|Q6AY56|TBA8_RAT  HVPRAVMVDLEPTVVDEVRAGTYRQLFHPEQLITGKEDAANNYARGHYTVGKESIDLVLD 120
                    ******:*******:**:* * ***************************:*** :* ***

SP|P68370|TBA1A_RAT RIRKLADQCTGLQGFLVFHSFGGGTGSGFTSLLMERLSVDYGKKSKLEFSIYPAPQVSTA 180
SP|Q6P9V9|TBA1B_RAT RIRKLADQCTGLQGFLVFHSFGGGTGSGFTSLLMERLSVDYGKKSKLEFSIYPAPQVSTA 180
SP|Q6AYZ1|TBA1C_RAT RIRKLADQCTGLQGFLVFHSFGGGTGSGFTSLLMERLSVDYGKKSKLEFSIYPAPQVSTA 180
SP|Q68FR8|TBA3_RAT  RIRKLADLCTGLQGFLIFHSFGGGTGSGFASLLMERLSVDYGKKSKLEFAIYPAPQVSTA 180
SP|Q5XIF6|TBA4A_RAT RIRKLSDQCTGLQGFLVFHSFGGGTGSGFTSLLMERLSVDYGKKSKLEFSIYPAPQVSTA 180
SP|Q6AY56|TBA8_RAT  RIRKLTDACSGLQGFLIFHSFGGGTGSGFTSLLMERLSLDYGKKSKLEFAIYPAPQVSTA 180
                    *****:* *:******:***********:*******:*********:*********

SP|P68370|TBA1A_RAT VVEPYNSILTTHTTLEHSDCAFMVDNEAIYDICRRNLDIERPTYTNLNRLIGQIVSSITA 240
SP|Q6P9V9|TBA1B_RAT VVEPYNSILTTHTTLEHSDCAFMVDNEAIYDICRRNLDIERPTYTNLNRLISQIVSSITA 240
SP|Q6AYZ1|TBA1C_RAT VVEPYNSILTTHTTLEHSDCAFMVDNEAIYDICRRNLDIERPTYTNLNRLISQIVSSITA 240
SP|Q68FR8|TBA3_RAT  VVEPYNSILTTHTTLEHSDCAFMVDNEAIYDICRRNLDIERPTYTNLNRLIGQIVSSITA 240
SP|Q5XIF6|TBA4A_RAT VVEPYNSILTTHTTLEHSDCAFMVDNEAIYDICRRNLDIERPTYTNLNRLISQIVSSITA 240
SP|Q6AY56|TBA8_RAT  VVEPYNSILTTHTTLEHSDCAFMVDNEAIYDICRRNLDIERPTYTNLNRLISQIVSSITA 240
                    ***************************************************.********

SP|P68370|TBA1A_RAT SLRFDGALNVDLTEFQTNLVPYPRIHFPLATYAPVISAEKAYHEQLSVAEITNACFEPAN 300
SP|Q6P9V9|TBA1B_RAT SLRFDGALNVDLTEFQTNLVPYPRIHFPLATYAPVISAEKAYHEQLSVAEITNACFEPAN 300
SP|Q6AYZ1|TBA1C_RAT SLRFDGALNVDLTEFQTNLVPYPRIHFPLATYAPVISAEKAYHEQLTVAEITNACFEPAN 300
SP|Q68FR8|TBA3_RAT  SLRFDGALNVDLTEFQTNLVPYPRIHFPLATYAPVISAEKAYHEQLSVAEITNACFEPAN 300
SP|Q5XIF6|TBA4A_RAT SLRFDGALNVDLTEFQTNLVPYPRIHFPLATYAPVISAEKAYHEQLSVAEITNACFEPAN 300
SP|Q6AY56|TBA8_RAT  SLRFDGALNVDLTEFQTNLVPYPRIHFPLVTYAPIVSAEKAYHEQLSVAEITSSCFEPNS 300
                    *****************************.****::**********:*****.:****  .

SP|P68370|TBA1A_RAT QMVKCDPRHGKYMACCLLYRGDVVPKDVNAAIATIKTKRTIQFVDWCPTGFKVGINYQPP 360
SP|Q6P9V9|TBA1B_RAT QMVKCDPRHGKYMACCLLYRGDVVPKDVNAAIATIKTKRSIQFVDWCPTGFKVGINYQPP 360
SP|Q6AYZ1|TBA1C_RAT QMVKCDPRHGKYMACCLLYRGDVVPKDVNAAIATIKTKRTIQFVDWCPTGFKVGINYQPP 360
SP|Q68FR8|TBA3_RAT  QMVKCDPRHGKYMACCMLYRGDVVPKDVNAAIATIKTKRTIQFVDWCPTGFKVGINYQPP 360
SP|Q5XIF6|TBA4A_RAT QMVKCDPRHGKYMACCLLYRGDVVPKDVNAAIAAIKTKRSIQFVDWCPTGFKVGINYQPP 360
SP|Q6AY56|TBA8_RAT  QMVKCDPRHGKYMACCMLYRGDVVPKDVNVAIAAIKTKRTIQFVDWCPTGFKVGINYQPP 360
                    ****************:************.***:*****:********************

SP|P68370|TBA1A_RAT TVVPGGDLAKVQRAVCMLSNTTAIAEAWARLDHKFDLMYAKRAFVHWYVGEGMEEGEFSE 420
SP|Q6P9V9|TBA1B_RAT TVVPGGDLAKVQRAVCMLSNTTAIAEAWARLDHKFDLMYAKRAFVHWYVGEGMEEGEFSE 420
SP|Q6AYZ1|TBA1C_RAT TVVPGGDLARVQRAVCMLSNTTAIAEAWARLDHKFDLMYAKRAFVHWYVGEGMEEGEFSE 420
SP|Q68FR8|TBA3_RAT  TVVPGGDLAKVQRAVCMLSNTTAIAEAWARLDHKFDLMYAKRAFVHWYVGEGMEEGEFSE 420
SP|Q5XIF6|TBA4A_RAT TVVPGGDLAKVQRAVCMLSNTTAIAEAWARLDHKFDLMYAKRAFVHWYVGEGMEEGEFSE 420
SP|Q6AY56|TBA8_RAT  TVVPGGDLAKVQRAVCMLSNTTAIAEAWARLDHKFDLMYAKRAFVHWYVGEGMEEGEFSE 420
                    *********:**************************************************

SP|P68370|TBA1A_RAT AREDMAALEKDYEEVGVDSVEGEGEEEGEEY 451
SP|Q6P9V9|TBA1B_RAT AREDMAALEKDYEEVGVDSVEGEGEEEGEEY 451
SP|Q6AYZ1|TBA1C_RAT AREDMAALEKDYEEVGADSAEGD--DEGEEY 449
SP|Q68FR8|TBA3_RAT  AREDLAALEKDYEEVGVDSVEAEAE-EGEEY 450
SP|Q5XIF6|TBA4A_RAT AREDMAALEKDYEEVGIDSYEDEDEGEE--- 448
SP|Q6AY56|TBA8_RAT  AREDLAALEKDYEEVGTDSFEEENEGEEF-- 449
                    ****:*********** ** * :   *
```

**Figure S3:** Sequence alignment of identified α-tubulin isoforms with modified residues labeled (**green** – 1 modification, **violet** – 2 modifications).

### β-TUBULIN

```
SP|P85108|TBB2A_RAT MREIVHIQAGQCGNQIGAKFWEVISDEHGIDPTGSYHGDSDLQLERINVYYNEAAGNKYV 60
SP|Q3KRE8|TBB2B_RAT MREIVHIQAGQCGNQIGAKFWEVISDEHGIDPTGSYHGDSDLQLERINVYYNEATGNKYV 60
SP|Q6P9T8|TBB4B_RAT MREIVHLQAGQCGNQIGAKFWEVISDEHGIDPTGTYHGDSDLQLERINVYYNEATGGKYV 60
SP|P69897|TBB5_RAT  MREIVHIQAGQCGNQIGAKFWEVISDEHGIDPTGTYHGDSDLQLDRISVYYNEATGGKYV 60
SP|Q4QRB4|TBB3_RAT  MREIVHIQAGQCGNQIGAKFWEVISDEHGIDPSGNYVGDSDLQLERISVYYNEASSHKYV 60
                    ******:*************************:*.* *******:**.******:. ***

SP|P85108|TBB2A_RAT PRAILVDLEPGTMDSVRSGPFGQIFRPDNFVFGQSGAGNNWAKGHYTEGAELVDSVLDVV 120
SP|Q3KRE8|TBB2B_RAT PRAILVDLEPGTMDSVRSGPFGQIFRPDNFVFGQSGAGNNWAKGHYTEGAELVDSVLDVV 120
SP|Q6P9T8|TBB4B_RAT PRAVLVDLEPGTMDSVRSGPFGQIFRPDNFVFGQSGAGNNWAKGHYTEGAELVDSVLDVV 120
SP|P69897|TBB5_RAT  PRAILVDLEPGTMDSVRSGPFGQIFRPDNFVFGQSGAGNNWAKGHYTEGAELVDSVLDVV 120
SP|Q4QRB4|TBB3_RAT  PRAILVDLEPGTMDSVRSGAFGHLFRPDNFIFGQSGAGNNWAKGHYTEGAELVDSVLDVV 120
                    ***:*************** **::******:*****************************

SP|P85108|TBB2A_RAT RKESESCDCLQGFQLTHSLGGGTGSGMGTLLISKIREEYPDRIMNTFSVMPSPKVSDTVV 180
SP|Q3KRE8|TBB2B_RAT RKESESCDCLQGFQLTHSLGGGTGSGMGTLLISKIREEYPDRIMNTFSVMPSPKVSDTVV 180
SP|Q6P9T8|TBB4B_RAT RKEAESCDCLQGFQLTHSLGGGTGSGMGTLLISKIREEYPDRIMNTFSVVPSPKVSDTVV 180
SP|P69897|TBB5_RAT  RKEAESCDCLQGFQLTHSLGGGTGSGMGTLLISKIREEYPDRIMNTFSVVPSPKVSDTVV 180
SP|Q4QRB4|TBB3_RAT  RKECENCDCLQGFQLTHSLGGGTGSGMGTLLISKVREEYPDRIMNTFSVVPSPKVSDTVV 180
                    ***.*.****************************:**************:**********

SP|P85108|TBB2A_RAT EPYNATLSVHQLVENTDETYSIDNEALYDICFRTLKLTTPTYGDLNHLVSATMSGVTTCL 240
SP|Q3KRE8|TBB2B_RAT EPYNATLSVHQLVENTDETYCIDNEALYDICFRTLKLTTPTYGDLNHLVSATMSGVTTCL 240
SP|Q6P9T8|TBB4B_RAT EPYNATLSVHQLVENTDETYCIDNEALYDICFRTLKLTTPTYGDLNHLVSATMSGVTACL 240
SP|P69897|TBB5_RAT  EPYNATLSVHQLVENTDETYCIDNEALYDICFRTLKLTTPTYGDLNHLVSATMSGVTTCL 240
SP|Q4QRB4|TBB3_RAT  EPYNATLSIHQLVENTDETYCIDNEALYDICFRTLKLATPTYGDLNHLVSATMSGVTTSL 240
                    ********:***********.****************:*******************:.*

SP|P85108|TBB2A_RAT RFPGQLNADLRKLAVNMVPFPRLHFFMPGFAPLTSRGSQQYRALTVPELTQQMFDSKNMM 300
SP|Q3KRE8|TBB2B_RAT RFPGQLNADLRKLAVNMVPFPRLHFFMPGFAPLTSRGSQQYRALTVPELTQQMFDSKNMM 300
SP|Q6P9T8|TBB4B_RAT RFPGQLNADLRKLAVNMVPFPRLHFFMPGFAPLTSRGSQQYRALTVPELTQQMFDAKNMM 300
SP|P69897|TBB5_RAT  RFPGQLNADLRKLAVNMVPFPRLHFFMPGFAPLTSRGSQQYRALTVPELTQQVFDAKNMM 300
SP|Q4QRB4|TBB3_RAT  RFPGQLNADLRKLAVNMVPFPRLHFFMPGFAPLTARGSQQYRALTVPELTQQMFDAKNMM 300
                    **********************************:*****************:**:****

SP|P85108|TBB2A_RAT AACDPRHGRYLTVAAIFRGRMSMKEVDEQMLNVQNKNSSYFVEWIPNNVKTAVCDIPPRG 360
SP|Q3KRE8|TBB2B_RAT AACDPRHGRYLTVAAIFRGRMSMKEVDEQMLNVQNKNSSYFVEWIPNNVKTAVCDIPPRG 360
SP|Q6P9T8|TBB4B_RAT AACDPRHGRYLTVAAVFRGRMSMKEVDEQMLNVQNKNSSYFVEWIPNNVKTAVCDIPPRG 360
SP|P69897|TBB5_RAT  AACDPRHGRYLTVAAVFRGRMSMKEVDEQMLNVQNKNSSYFVEWIPNNVKTAVCDIPPRG 360
SP|Q4QRB4|TBB3_RAT  AACDPRHGRYLTVATVFRGRMSMKEVDEQMLAIQSKNSSYFVEWIPNNVKVAVCDIPPRG 360
                    **************::***************  :*.**************.*********

SP|P85108|TBB2A_RAT LKMSATFIGNSTAIQELFKRISEQFTAMFRRKAFLHWYTGEGMDEMEFTEAESNMNDLVS 420
SP|Q3KRE8|TBB2B_RAT LKMSATFIGNSTAIQELFKRISEQFTAMFRRKAFLHWYTGEGMDEMEFTEAESNMNDLVS 420
SP|Q6P9T8|TBB4B_RAT LKMSATFIGNSTAIQELFKRISEQFTAMFRRKAFLHWYTGEGMDEMEFTEAESNMNDLVS 420
SP|P69897|TBB5_RAT  LKMAVTFIGNSTAIQELFKRISEQFTAMFRRKAFLHWYTGEGMDEMEFTEAESNMNDLVS 420
SP|Q4QRB4|TBB3_RAT  LKMSSTFIGNSTAIQELFKRISEQFTAMFRRKAFLHWYTGEGMDEMEFTEAESNMNDLVS 420
                    ***: *******************************************************

SP|P85108|TBB2A_RAT EYQQYQDATADEQGEFEEEEGEDEA----- 445
SP|Q3KRE8|TBB2B_RAT EYQQYQDATADEQGEFEEEEGEDEA----- 445
SP|Q6P9T8|TBB4B_RAT EYQQYQDATAEEEGEFEEEAEEEVA----- 445
SP|P69897|TBB5_RAT  EYQQYQDATAEEEEDFGEEAEEEA------ 444
SP|Q4QRB4|TBB3_RAT  EYQQYQDATAEEEGEMYEDDDEESEAQGPK 450
                    **********:*: :: *:  *:
```

**Figure S4:** Sequence alignment of identified β-tubulin isoforms with modified residues labeled (**green** – 1 modification, **violet** – 2 modifications, **red** – 3 modifications, **orange** – 4 modifications).

**Vimentin**

sp|P31000|VIME_RAT
MSTRSVSSSSYRRMFGGSGTSSRPSSNRSYVTTSTRTYSLGSALRPSTSRSLYSSSPGGA
YVTRSSAVRLRSSMPGVRLLQDSVDFSLADAINTEFKNTRTNEKVELQELNDRFANYIDK
VRFLEQQNKILLAELEQLKGQGKSRLGDLYEEEMRELRRQVDQLTNDKARVEVERDNLAE
DIMRLREKLQEEMLQREEAESTLQSFRQDVDNASLARLDLERKVESLQEEIAFLKKLHDE
EIQELQAQIQEQHVQIDVDVSKPDLTAALRDVRQQYESVAAKNLQEAEEWYKSKFADLSE
AANRNNDALRQAKQESNEYRRQVQSLTCEVDALKGTNESLERQMREMEENFALEAANYQD
TIGRLQDEIQNMKEEMARHLREYQDLLNVKMALDIEIATYRKLLEGEESRISLPLPNFSS
LNLRETNLESLPLVDTHSKRTLLIKTVETRDGQVINETSQHHDDLE

**Figure S5:** Amino acid sequence of vimentin with modified residues labeled (**green** – 1 modification, **violet** – 2 modifications, **red** – 3 modifications, **orange** – 4 modifications, **dark red** – 5 modifications, **pink** – 6 modifications, **turquoise** – 7 modifications).

### Actin

```
SP|P60711|ACTB_RAT --MDDDIAALVVDNGSGMCKAGFAGDDAPRAVFPSIVGRPRHQGVMVGMGQKDSYVGDEA 58
SP|P63259|ACTG_RAT --MEEEIAALVIDNGSGMCKAGFAGDDAPRAVFPSIVGRPRHQGVMVGMGQKDSYVGDEA 58
SP|P68035|ACTC_RAT MCDDEETTALVCDNGSGLVKAGFAGDDAPRAVFPSIVGRPRHQGVMVGMGQKDSYVGDEA 60
SP|P62738|ACTA_RAT MCEEEDSTALVCDNGSGLCKAGFAGDDAPRAVFPSIVGRPRHQGVMVGMGQKDSYVGDEA 60
SP|P63269|ACTH_RAT MCEE-ETTALVCDNGSGLCKAGFAGDDAPRAVFPSIVGRPRHQGVMVGMGQKDSYVGDEA 59
SP|P68136|ACTS_RAT MCDEDETTALVCDNGSGLVKAGFAGDDAPRAVFPSIVGRPRHQGVMVGMGQKDSYVGDEA 60
                      : : :*** *****: *****************************************

SP|P60711|ACTB_RAT QSKRGILTLKYPIEHGIVTNWDDMEKIWHHTFYNELRVAPEEHPVLLTEAPLNPKANREK 118
SP|P63259|ACTG_RAT QSKRGILTLKYPIEHGIVTNWDDMEKIWHHTFYNELRVAPEEHPVLLTEAPLNPKANREK 118
SP|P68035|ACTC_RAT QSKRGILTLKYPIEHGIITNWDDMEKIWHHTFYNELRVAPEEHPTLLTEAPLNPKANREK 120
SP|P62738|ACTA_RAT QSKRGILTLKYPIEHGIITNWDDMEKIWHHSFYNELRVAPEEHPTLLTEAPLNPKANREK 120
SP|P63269|ACTH_RAT QSKRGILTLKYPIEHGIITNWDDMEKIWHHSFYNELRVAPEEHPTLLTEAPLNPKANREK 119
SP|P68136|ACTS_RAT QSKRGILTLKYPIEHGIITNWDDMEKIWHHTFYNELRVAPEEHPTLLTEAPLNPKANREK 120
                   *****************:************:*************.***************

SP|P60711|ACTB_RAT MTQIMFETFNTPAMYVAIQAVLSLYASGRTTGIVMDSGDGVTHTVPIYEGYALPHAILRL 178
SP|P63259|ACTG_RAT MTQIMFETFNTPAMYVAIQAVLSLYASGRTTGIVMDSGDGVTHTVPIYEGYALPHAILRL 178
SP|P68035|ACTC_RAT MTQIMFETFNVPAMYVAIQAVLSLYASGRTTGIVLDSGDGVTHNVPIYEGYALPHAIMRL 180
SP|P62738|ACTA_RAT MTQIMFETFNVPAMYVAIQAVLSLYASGRTTGIVLDSGDGVTHNVPIYEGYALPHAIMRL 180
SP|P63269|ACTH_RAT MTQIMFETFNVPAMYVAIQAVLSLYASGRTTGIVLDSGDGVTHNVPIYEGYALPHAIMRL 179
SP|P68136|ACTS_RAT MTQIMFETFNVPAMYVAIQAVLSLYASGRTTGIVLDSGDGVTHNVPIYEGYALPHAIMRL 180
                   **********.***********************:********.*************:**

SP|P60711|ACTB_RAT DLAGRDLTDYLMKILTERGYSFTTTAEREIVRDIKEKLCYVALDFEQEMATAASSSSLEK 238
SP|P63259|ACTG_RAT DLAGRDLTDYLMKILTERGYSFTTTAEREIVRDIKEKLCYVALDFEQEMATAASSSSLEK 238
SP|P68035|ACTC_RAT DLAGRDLTDYLMKILTERGYSFVTTAEREIVRDIKEKLCYVALDFENEMATAASSSSLEK 240
SP|P62738|ACTA_RAT DLAGRDLTDYLMKILTERGYSFVTTAEREIVRDIKEKLCYVALDFENEMATAASSSSLEK 240
SP|P63269|ACTH_RAT DLAGRDLTDYLMKILTERGYSFVTTAEREIVRDIKEKLCYVALDFENEMATAASSSSLEK 239
SP|P68136|ACTS_RAT DLAGRDLTDYLMKILTERGYSFVTTAEREIVRDIKEKLCYVALDFENEMATAASSSSLEK 240
                   **********************.***********************:*************

SP|P60711|ACTB_RAT SYELPDGQVITIGNERFRCPEALFQPSFLGMESCGIHETTFNSIMKCDVDIRKDLYANTV 298
SP|P63259|ACTG_RAT SYELPDGQVITIGNERFRCPEALFQPSFLGMESCGIHETTFNSIMKCDVDIRKDLYANTV 298
SP|P68035|ACTC_RAT SYELPDGQVITIGNERFRCPETLFQPSFIGMESAGIHETTYNSIMKCDIDIRKDLYANNV 300
SP|P62738|ACTA_RAT SYELPDGQVITIGNERFRCPETLFQPSFIGMESAGIHETTYNSIMKCDIDIRKDLYANNV 300
SP|P63269|ACTH_RAT SYELPDGQVITIGNERFRCPETLFQPSFIGMESAGIHETTYNSIMKCDIDIRKDLYANNV 299
SP|P68136|ACTS_RAT SYELPDGQVITIGNERFRCPETLFQPSFIGMESAGIHETTYNSIMKCDIDIRKDLYANNV 300
                   *********************:******:****.******:*******:*********.*

SP|P60711|ACTB_RAT LSGGTTMYPGIADRMQKEITALAPSTMKIKIIAPPERKYSVWIGGSILASLSTFQQMWIS 358
SP|P63259|ACTG_RAT LSGGTTMYPGIADRMQKEITALAPSTMKIKIIAPPERKYSVWIGGSILASLSTFQQMWIS 358
SP|P68035|ACTC_RAT LSGGTTMYPGIADRMQKEITALAPSTMKIKIIAPPERKYSVWIGGSILASLSTFQQMWIS 360
SP|P62738|ACTA_RAT LSGGTTMYPGIADRMQKEITALAPSTMKIKIIAPPERKYSVWIGGSILASLSTFQQMWIS 360
SP|P63269|ACTH_RAT LSGGTTMYPGIADRMQKEITALAPSTMKIKIIAPPERKYSVWIGGSILASLSTFQQMWIS 359
SP|P68136|ACTS_RAT MSGGTTMYPGIADRMQKEITALAPSTMKIKIIAPPERKYSVWIGGSILASLSTFQQMWIT 360
                   :**********************************************************:

SP|P60711|ACTB_RAT KQEYDESGPSIVHRKCF 375
SP|P63259|ACTG_RAT KQEYDESGPSIVHRKCF 375
SP|P68035|ACTC_RAT KQEYDEAGPSIVHRKCF 377
SP|P62738|ACTA_RAT KQEYDEAGPSIVHRKCF 377
SP|P63269|ACTH_RAT KPEYDEAGPSIVHRKCF 376
SP|P68136|ACTS_RAT KQEYDEAGPSIVHRKCF 377
                   * ****:**********
```

**Figure S6:** Sequence alignment of identified actin isoforms with modified residues labeled (**green** – 1 modification, **violet** – 2 modifications, **red** – 3 modifications, **orange** – 4 modifications, **dark red** – 5 modifications, **pink** – 6 modifications).

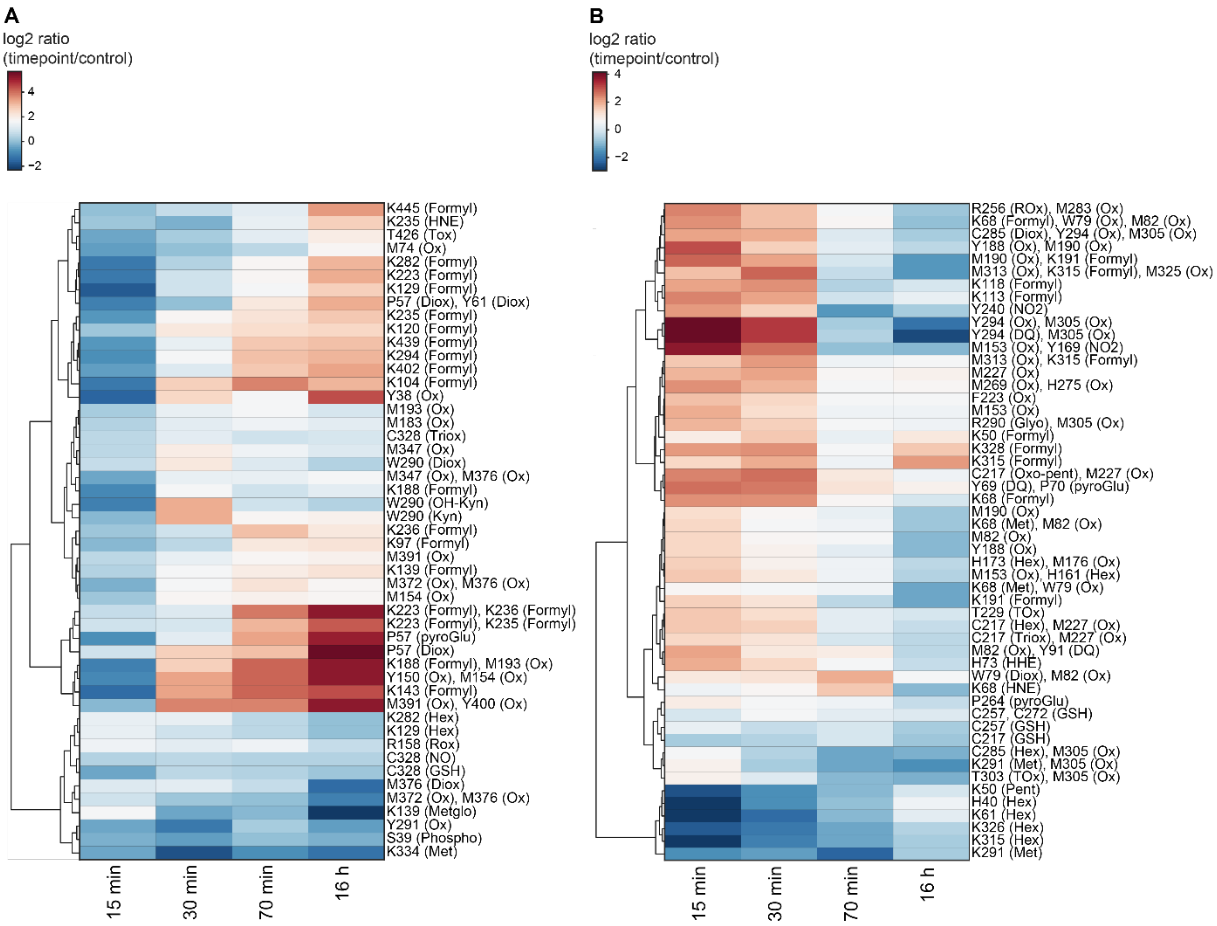


**Figure S7**: Hierarchical clustering of modification sites in (A) vimentin and (B) actin based on the relatively quantified corresponding tryptic peptides. Log2-transformed ratios of timepoints relative to control are depicted. DQ – dopaquinone, Diox – dioxidation, Formyl – formylation, Glyo – glyoxal, GSH – glutathionylation, Hex – hexenal, HHE – hydroxyhexenal, HNE – hydroxynonenal, Kyn – kynurenine, Metglo – methylglyoxal, Met – methylation, NO – nitrosation or nitrosylation, NO2 – nitration, OH-Kyn – hydroxykynurenine, Ox – oxidation, Oxo-pent – oxo-pentanal, Pent – pentanal, Phospho – phosphorylation, pyroGlu – pyroglutamic acid, Rox – glutamic semialdehyde, Tox – 2-Amino-3-ketobutyric acid, Triox – trioxidation. Hierarchical clustering was created using Instant Clue (version 0.5.3) (4).

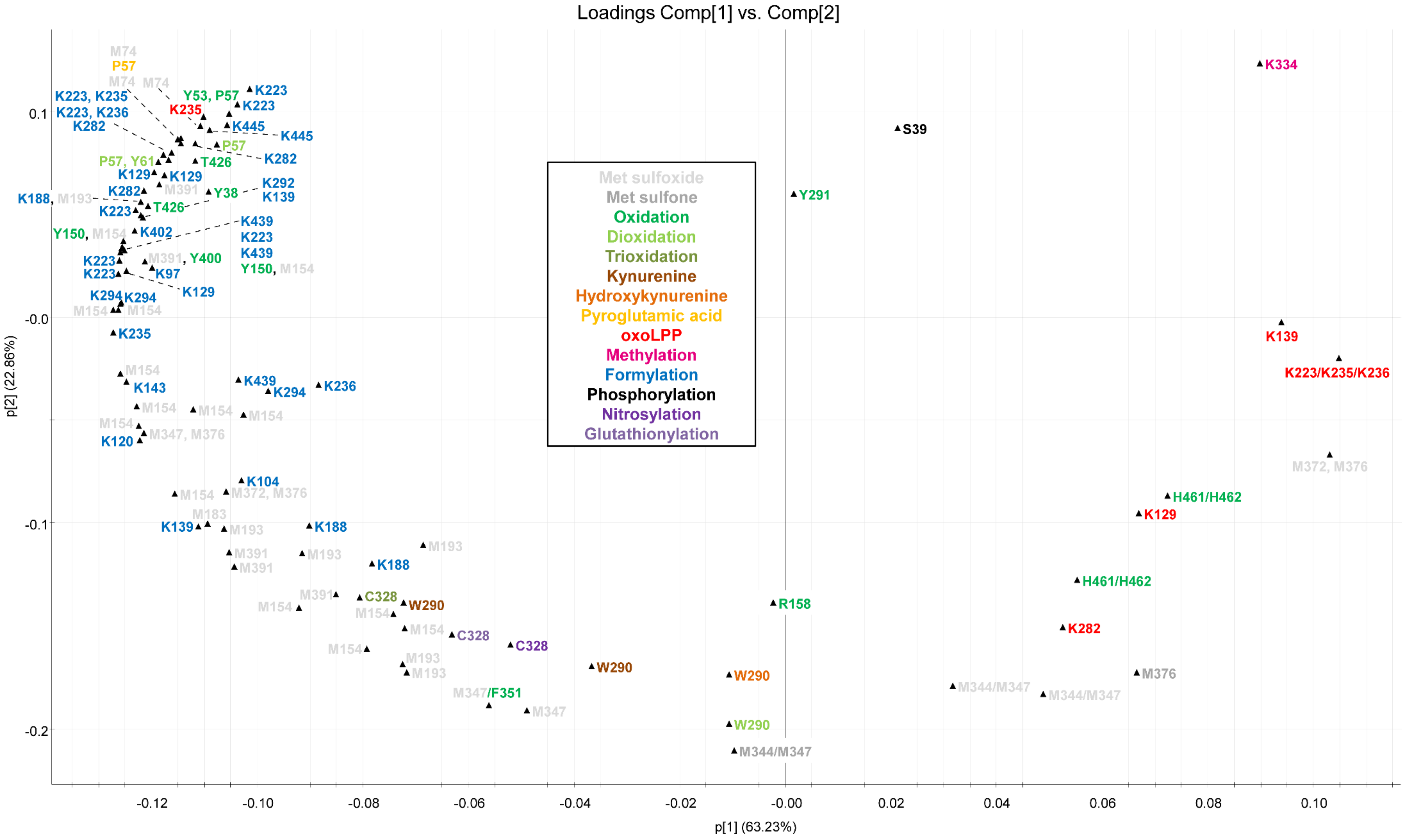


**Figure S8**: PCA loadings plot of modification sites in vimentin based on the relatively quantified corresponding tryptic peptides.

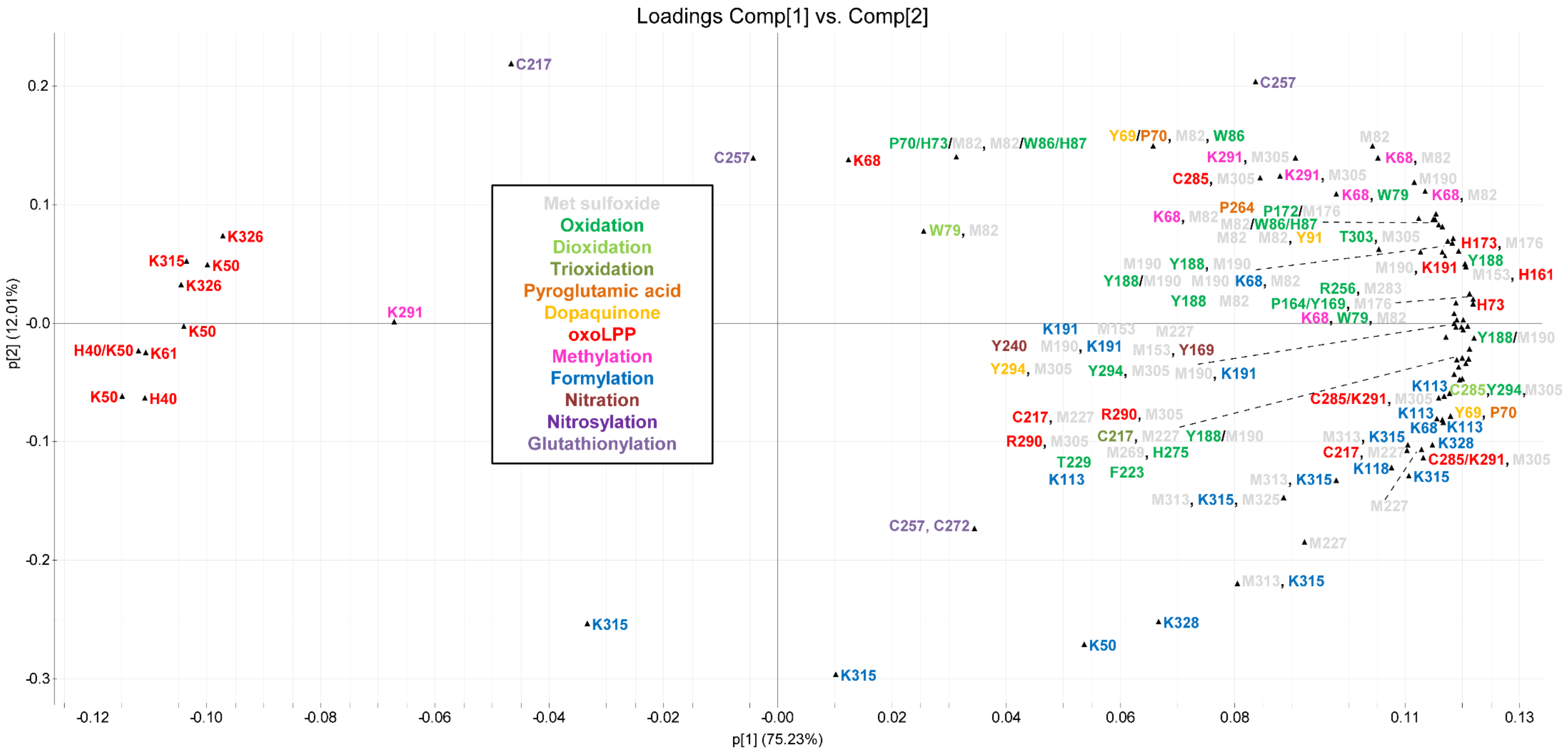


**Figure S9**: PCA loadings plot of modification sites in actin based on the relatively quantified corresponding tryptic peptides.
